# Frequent EPHA2 receptor mutations in cholangiocarcinoma disrupt receptor forward signaling supporting a tumor suppressor role

**DOI:** 10.1101/2025.02.25.640198

**Authors:** Evodie Koutouan, Ayano Niibe, Jack W. Sample, Danielle Carlson, Rondell Graham, Rory L. Smoot, Elena B. Pasquale

**Affiliations:** Cancer Center, Sanford Burnham Prebys Medical Discovery Institute, La Jolla, California 92037, USA; Division of Hepatobiliary and Pancreas Surgery & Department of Biochemistry and Molecular Biology, Mayo Clinic, Rochester, Minnesota 55905, USA; Department of Laboratory Medicine and Pathology, Mayo Clinic, Rochester, Minnesota 55905, USA

**Keywords:** receptor tyrosine kinase, phosphorylation, cancer driver mutation, tumor initiation

## Abstract

EPHA2 is a receptor tyrosine kinase highly expressed in many cancers. By analyzing cancer patient databases for mutations in the EPHA2 coding sequence, we found that cholangiocarcinoma (a hepatobiliary cancer with dismal prognosis) exhibits a uniquely high incidence of EPHA2 mutations. To define the functional significance of these mutations, we generated representative EPHA2 mutants and monitored major receptor autophosphorylation sites as indicative of kinase activity-dependent signal transduction (known as forward signaling). We found that missense mutations in the ligand-binding domain abrogate ligand binding and ligand-induced EPHA2 tyrosine phosphorylation, while most missense mutations in the kinase domain abrogate kinase activity. We detected less pronounced effects of missense mutations in other domains, which vary depending on the phosphosite, suggesting that these EPHA2 mutations might differentially affect (or bias) different downstream signaling pathways. Other EPHA2 mutations introduce early stop codons and encode receptor truncated forms lacking all or part of the kinase domain. We also found that an EPHA2 secreted truncated form, a transmembrane truncated form, and a full-length kinase inactive mutant can inhibit tyrosine phosphorylation of co-expressed EPHA2 wild-type. Taken together, these data suggest that mutations interfering with EPHA2 forward signaling facilitate cholangiocarcinoma development. We indeed obtained evidence that an EPHA2 kinase inactive mutant, but not EPHA2 wild-type, can induce proliferative masses consistent with well differentiated cholangiocarcinoma in a validated mouse model of cholangiocarcinogenesis. Taken together, our findings suggest that EPHA2 is a driver gene in cholangiocarcinoma and that its forward signaling has tumor suppressor activity.

## INTRODUCTION

Cholangiocarcinoma (CCA) is a cancer arising from the biliary epithelia (1–3). The incidence of this cancer is increasing and because of its clinically silent presentation, patients are often diagnosed at advanced disease stages, when curative-intent treatments are not possible. Palliative approaches have had very minimal impact on patient survival, which remains at 7-20% overall (4–6). The current standard of care first-line systemic therapy is a combination of gemcitabine, cisplatin, and either durvalumab or pembrolizumab (7–10). This triplet therapy provides median survival of just over one year and compared to the doublet regimen of gemcitabine and cisplatin improves median survival by only about one month. Similarly, second line treatments with targeted agents in patients with tumors harboring FGFR2 fusions, IDH1 mutations or other targetable mutations have limited overall response rates (typically less than 40%) and durations of response that are measured in months (2, 3, 11–16). As a consequence, despite being a rare cancer, CCA is responsible for ∼2% of cancer related deaths worldwide (6). Since currently approved treatments have provided small incremental benefits to patients, new insights into the molecular alterations associated with CCA development and progression are needed. Of special interest are molecular changes that have implications for CCA development, since earlier interception is a promising strategy to improve overall survival, given the limited treatment options for CCA.

Clinical risk factors associated with CCA development have been identified. However, specific molecular alterations and mechanisms underlying these changes are unknown. For example, inflammatory conditions such as primary sclerosing cholangitis or liver fluke infection are associated with markedly increased risk of CCA development (1, 3, 6, 17, 18). While public health initiatives have impacted the incidence of liver fluke infection as a risk factor, patients with primary sclerosing cholangitis are at the highest risk (with ∼1,000-fold higher incidence than the general population). However, surveillance based on imaging and endoscopic sampling have not impacted the rate of CCA development. Current molecular diagnostics, which include fluorescence *in situ* hybridization probes for genes including MCL1, EGFR, MYC, and CDKN2A, are utilized for cancer diagnosis in cytology specimens but do not seem useful for risk stratification (19–21).

Genetically engineered mouse models of cholangiocarcinogenesis most commonly involve expression of known oncogenes and/or elimination of tumor suppressor genes (such as *KRAS* and *TP53*, respectively) or phenocopy signaling changes without specifically genocopying the mutations found in the disease. For example, signaling by the oncogenic serine/threonine kinase AKT is commonly elevated in CCA tumors and constitutively active myristoylated AKT (myrAKT) is expressed as a permissive alteration (17, 22, 23). Activity of the transcriptional coactivator YAP1 is also high in most human CCA tumors, although not by direct mutation, and the combination of myrAKT and activated (S127A mutant) YAP1 induces CCA in mouse models (17, 22). On the other hand, the tumor suppressor FBXW7 has low expression in a majority of CCA tumors, and myrAKT in combination with a dominant negative form of Fbxw7 (Fbxw7ΔF) induces CCA in mouse models (22, 23). Thus, the majority of CCA mouse models functionally mimic molecular alterations found in human CCA tumors but do not accurately recapitulate them, since the tumor promoting activity of individual alterations is not fully defined. Overall, the mechanisms leading to malignant transformation of the biliary epithelium and subsequent tumor progression remain unknown.

Therefore, new molecular insights into the development and progression of CCA are needed to provide meaningful advances for patients in the areas of risk mitigation, diagnosis, and therapeutics.

By analyzing multiple human CCA databases available in cBioPortal (cbioportal.org), we found that the ephrin receptor EPHA2 exhibits a uniquely high incidence of coding sequence mutations in CCA compared to other human cancers (cbioportal.org). EPHA2 is a member of the Eph receptor tyrosine kinase family, which is highly expressed in most cancer types (24–27). The Eph receptor family includes 9 EphA and 5 EphB receptors, which preferentially bind the glycosylphosphatidylinositol-linked ephrinA ligands or the transmembrane ephrinB ligands, respectively. The extracellular portion of the Eph receptors contains an N-terminal ligand-binding domain, a cysteine-rich region that includes a Sushi domain and an epidermal growth factor (EGF)- like domain, and two fibronectin type III domains. The intracellular portion of the Eph receptors contains a juxtamembrane segment, a tyrosine kinase domain, a sterile-alpha-motif (SAM) domain, and a C-terminal PDZ domain-binding motif (Fig. 1).

**Figure 1.**
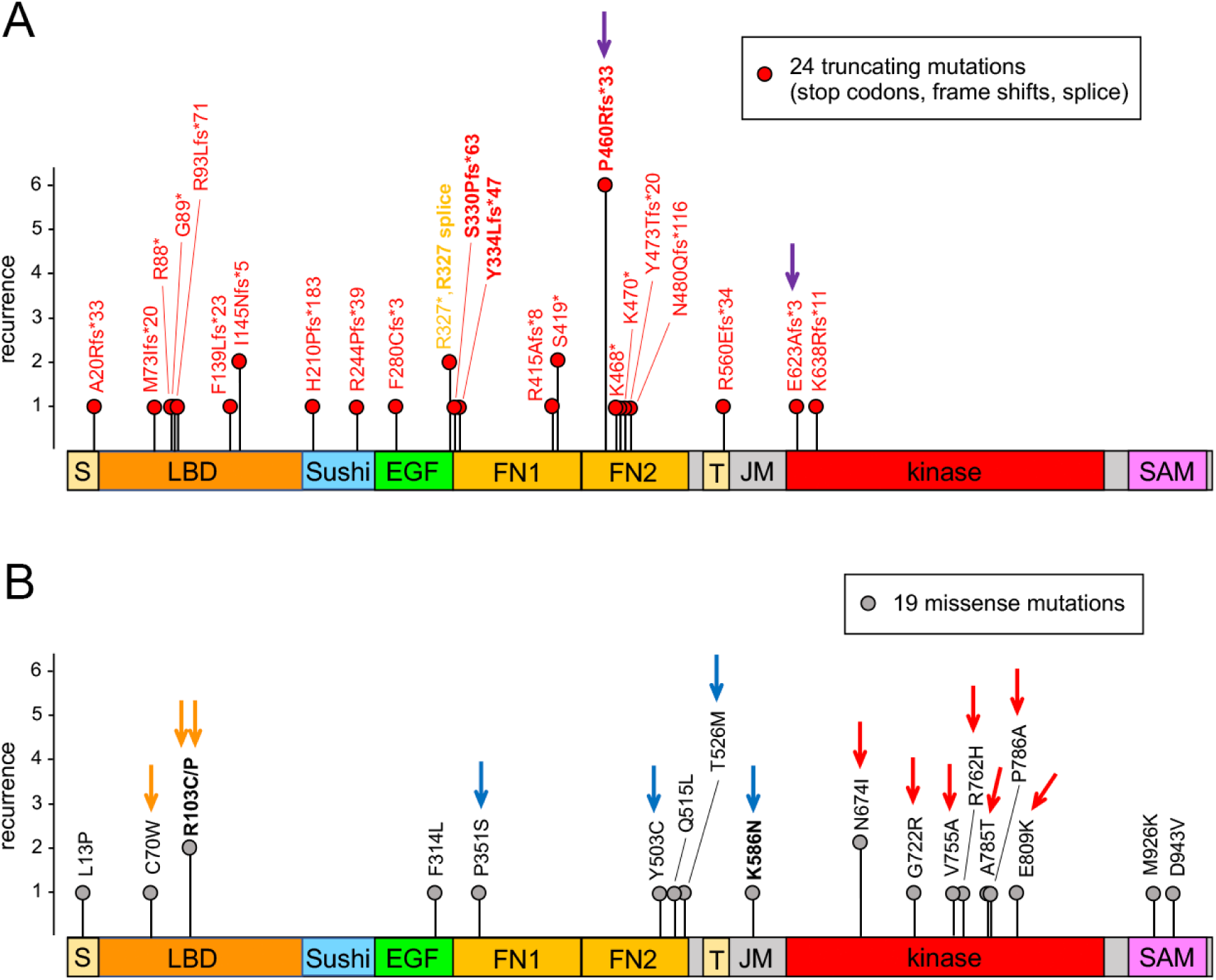
EPHA2 mutations in patient tumors from 6 cholangiocarcinoma studies. EPHA2 mutations identified in 611 tumors profiled for EPHA2 mutations in 6 cholangiocarcinoma studies (Fig. S1B; cbioportal.org) include nonsense mutations (*, red font), frame-shift mutations (fs, red font) and splice site mutations (orange font) (**A**) and missense mutations (**B**) (see also Table S1). The height of the bars indicates the number of tumors with a particular mutation (recurrence). The recurrent mutations that were also identified in the biliary tract tumors in the China Pan-cancer study (Fig. S2B,C; Table S2) are in bold. Arrows of different colors mark the different groups of mutations studied (purple, truncating mutations; orange, LBD mutations; blue, mutations in the fibronectin domains and juxtamembrane segment; red, kinase domain mutations). The EPHA2 domains are indicated: SP, signal peptide; LBD, ligand-binding domain; EGF, EGF-like domain; Sushi domain; FN, fibronectin type III domain; TM, transmembrane helix; JM, juxtamembrane segment; kinase, kinase domain; SAM, sterile alpha motif domain.

Interaction of the Eph receptors with ephrin ligands, which typically occurs at sites of cell-cell contact, leads to bidirectional signaling (25, 26, 28). This includes forward signaling, which depends on receptor tyrosine kinase activity, and reverse signaling, occurring in the cells expressing the ephrin ligand. Forward signaling is similar to signaling by other receptor tyrosine kinases, and it is therefore also known as “canonical” signaling (27). In forward signaling, the binding of ephrin ligands induces Eph receptor oligomerization, which brings receptor molecules into close proximity enabling their mutual cross-phosphorylation on tyrosine residues (25–29). Phosphorylation of two conserved tyrosines in the juxtamembrane segment (Y588 and Y594 in EPHA2) and a conserved tyrosine in the activation loop of the kinase domain (Y772 in EPHA2) promotes Eph receptor tyrosine kinase activity. These and other tyrosine phosphorylated motifs mediate the binding of downstream signaling effectors that contain SH2 or phosphotyrosine-binding (PTB) domains, leading to modulation of downstream signaling networks. Signaling cascades triggered by the Eph receptors also involve proteins that bind to the C-terminal PDZ domain-binding motif or to other Eph receptor intracellular regions as well as signaling proteins whose activity is regulated by the activated Eph receptors through tyrosine phosphorylation.

The EPHA2 receptor (originally named ECK for epithelial cell kinase) is predominantly expressed in epithelial cells, where together with ephrinA ligands it regulates cell adhesion, movement and communication and plays a role in maintaining epithelial homeostasis (30–35). Dysregulation of EPHA2 function has also been implicated in many diseases, including inflammation, pathological angiogenesis, psoriasis, cataracts, parasite infections, and cancer (24, 25, 27, 36–41). EPHA2 is widely upregulated in many tumor types, but its forward signaling activity is often low in cancer cells (24, 42, 43). This is consistent with many reports showing that EPHA2 forward signaling inhibits oncogenic signaling networks, although tumor promoting effects of EPHA2 forward signaling have also been reported (24, 25, 27). The role of EPHA2 forward signaling in cancer is incompletely understood; in particular, little is known about the role of cancer mutations that affect EPHA2 signaling on tumor development and progression. Therefore, we sought to investigate how the many mutations identified in CCA affect EPHA2 forward signaling and whether they may play a role in tumor initiation. Taken together, our findings show that the majority of the mutations in biliary tract tumors inactivate EPHA2 forward signaling and may play a role in cholangiocarcinogenesis.

## RESULTS

### EPHA2 mutations are common in human cholangiocarcinoma tumors

By surveying Eph receptor mutations identified in 6 CCA studies profiled for mutations from the cBioPortal database (cbioportal.org), we found that EPHA2 is by far the most frequently mutated of the 14 Eph receptor family members (Fig. S1A,B). Consistent with this, two different platforms designed to predict cancer driver genes highlight a potential role of EPHA2 as a driver specifically in CCA and not any other tumor types (44, 45) (intogen.org). In addition, other CCA studies have identified EPHA2 as a significantly mutated gene (1, 46, 47). Of the CCA tumors examined for EPHA2 mutations in the 6 CCA studies, 8% were found to harbor EPHA2 mutations (Fig. S1B; Table S1). A similar pattern of mutations is also apparent in the more recent “China Pan-cancer study” (48) (cbioportal.org), which profiles EPHA2 mutations in biliary tract tumors (including intrahepatic and extrahepatic CCAs and gallbladder tumors) and was published while our work was in progress (Fig. S2; Table S2).

Overall, the majority (72%) of the EPHA2 mutations in biliary tract tumors are stop codons or frameshift mutations that are predicted to encode truncated forms of EPHA2 lacking the SAM domain and the entire kinase domain or a portion of the kinase domain (Figs. 1A and S2B). In some cases, these truncated forms are short N-terminal fragments not including any intact domain.

Furthermore, a few mutations also involve splice sites. The EPHA2 missense mutations are spread throughout the domains of the receptor (Figs. 1B and S2C), and many are predicted to strongly affect function (Tables S1 and S2). Finally, among the 8 biliary tract cell lines included in the Cancer Cell Line Encyclopedia (CCLE), the SNU1079 cell line harbors the EPHA2 T614P kinase domain mutation (cbioportal.org) (not shown). Consistent with a role of EPHA2 as a tumor suppressor, analysis of the 36 TCGA CCA tumors with gene mutation information shows that EPHA2 mutations occur in tumors also harboring putative EPHA2 heterozygous loss (cbioportal.org; Fig. S1C) (49). In fact, most (80%) of the TCGA CCA tumors exhibit putative EPHA2 heterozygous loss or homozygous deletion. The SNU245 CCLE cholangiocarcinoma cell line also harbors EPHA2 homozygous deletion (cbioportal.org) (not shown) To experimentally characterize the effects of the mutations on EPHA2 ligand binding and tyrosine phosphorylation (indicative of kinase activity (50)), we generated lentiviral constructs encoding 16 representative EPHA2 mutants (arrows in Figs. 1 and 2B,C) with an N-terminal FLAG tag and used them for transient or stable transfection of HEK293 cells.

**Figure 2.**
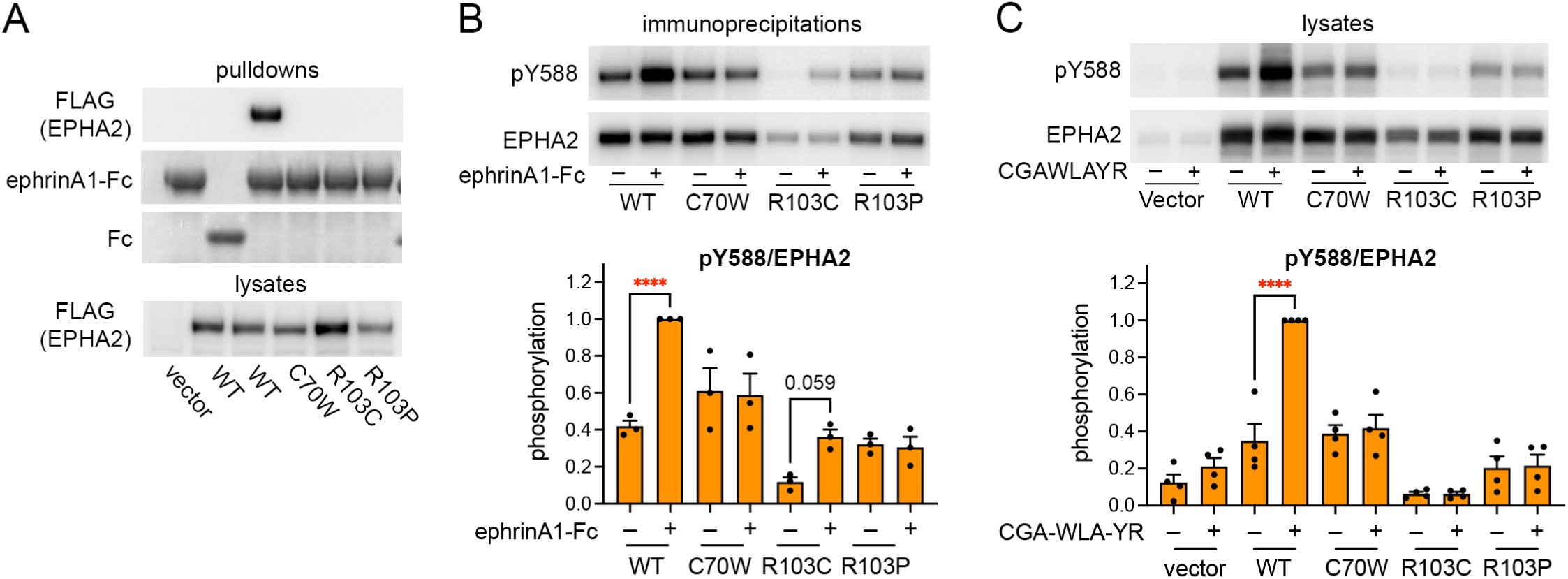
Missense mutations in the ligand-binding domain abrogate EPHA2 ligand binding. (**A**) EphrinA1-Fc and control Fc pulldowns from HEK293 cells transiently transfected with FLAG- EPHA2 WT and the indicated FLAG-EPHA2 LBD mutants were probed by immunoblotting with an anti-FLAG antibody to detect EPHA2 while the amido black protein stain was used to detect ephrinA1-Fc and Fc. Lysates were also probed for the FLAG epitope to detect EPHA2. (**B,C**) HEK293 cells stably expressing FLAG-EPHA2 WT and the indicated FLAG-EPHA2 LBD mutants were treated for 15 min with 0.25 ug/ml of the ephrinA1-Fc ligand followed by immunoprecipitation with an anti-FLAG antibody (B) or with 4 nM of the EPHA2-specific dimeric peptide agonist CGA-WLA-YR and lysed (C). Immunoprecipitates (B) and cell lysates (C) were probed for EPHA2 phosphorylation on tyrosine 588 (pY588) and for EPHA2. The bars in the graphs show averages ± SE for pY558/EPHA2 normalized to pY558/EPHA2 for ligand-stimulated EPHA2 WT. The individual measurements (from 3 experiments in B and 4 experiments in C) are shown as black dots. The asterisks indicate the significance of the difference between ligand treatment and no treatment, calculated using two-way ANOVA followed by the SV ıdák’s multiple comparisons test (****, p < 0.0001). The close to significant p-value for the R103C mutant is also shown.

### Missense mutations in the EPHA2 ligand-binding domain abrogate ligand binding

To investigate the impact of missense mutations in the EPHA2 LBD on receptor signaling, we generated the recurrent EPHA2 C70W, R103C, and R103P mutants identified in tumor samples of patients with biliary tract cancer (Figs. 1B and S2C). To assess the impact of the mutations on EPHA2 ligand binding ability, we transiently transfected HEK293 cells with EPHA2 wild-type (WT) and the three mutants. Cell lysates were used for pulldown assays using the engineered EPHA2 ligand ephrinA1 fused to the Fc portion of an antibody (ephrinA1-Fc) and immobilized on protein G beads. By immunoblotting the pulled down proteins associated with ephrinA1-Fc with a FLAG antibody, we detected EPHA2 WT but not the C70W, R103C, and R103P mutants (Fig. 2A), indicating that the mutations all drastically impair the ligand binding ability of EPHA2.

Binding of activating ligands to the EPHA2 receptor induces receptor cross-phosphorylation on tyrosine residues and downstream signaling (27). Detection of EPHA2 tyrosine phosphorylation with phosphospecific antibodies is an indirect but likely more sensitive method to detect low levels of residual ligand binding to EPHA2 as compared to ephrinA1-Fc pulldown. To examine the extent of signaling impairment caused by the LBD mutations, we treated stably transfected HEK293 cells with two ligands that activate EPHA2 forward signaling. We used the promiscuous ephrinA1-Fc ligand, which binds to all EphA receptors, and the CGA-WLA-YR peptide dimer, which is an engineered ligand that binds specifically to the ephrin-binding pocket of EPHA2 and not any other Eph receptor (50). We measured the effects of the ligands on EPHA2 phosphorylation on Y588 in the EPHA2 juxtamembrane region, which is a major autophosphorylation site that promotes EPHA2 kinase activity as well as interaction with signaling effectors containing SH2 domains (27). As expected, both ligands increased Y588 phosphorylation of EPHA2 WT (Fig. 2B,C). In contrast, none of the three mutants showed significantly increased Y588 phosphorylation in response to the two ligands. However, we noticed that the promiscuous ephrinA1-Fc, but not the EPHA2-specific peptide ligand, consistently induces low but detectable EPHA2 R103C phosphorylation (Fig. 2B,C). This suggests that an endogenous EphA receptor expressed in HEK293 cells, such as EPHA4 (our unpublished data), may be activated by ephrinA1-Fc and cross-phosphorylate the transfected EPHA2. This perhaps suggests that the R103C mutation favors association with other EphA receptors. Overall, these results show that all three mutations in the EPHA2 LBD drastically impair the ability of EPHA2 to bind ligands and its ligand-induced kinase-dependent signaling.

### Missense mutations in the EPHA2 kinase domain impair kinase activity

EPHA2 spontaneously oligomerizes when expressed at high levels in transiently transfected HEK293 cells, leading to cross-phosphorylation on multiple tyrosine residues, including not only Y588 but also Y594 (also in the juxtamembrane region) and Y772 (in the activation loop of the kinase domain). These three major autophosphorylation sites function in concert to promote EPHA2 kinase activity and downstream forward signaling (27). We investigated the impact on EPHA2 tyrosine phosphorylation of seven EPHA2 kinase domain mutations identified in CCA tumors: N674I (recurrent in two tumors), G722R, V755A, R762H, A785T, P786A, and E809K (Fig. 1B). By monitoring phosphorylation on specific tyrosines (Y588, Y594, and Y772) as well as overall tyrosine phosphorylation (pTyr), we found that six of the mutations essentially abolish EPHA2 tyrosine phosphorylation while the R762H mutation significantly reduces it (Fig. 3). Thus, a number of EPHA2 kinase domain missense mutations in CCA tumors strongly inhibit EPHA2 kinase activity and forward signaling.

**Figure 3.**
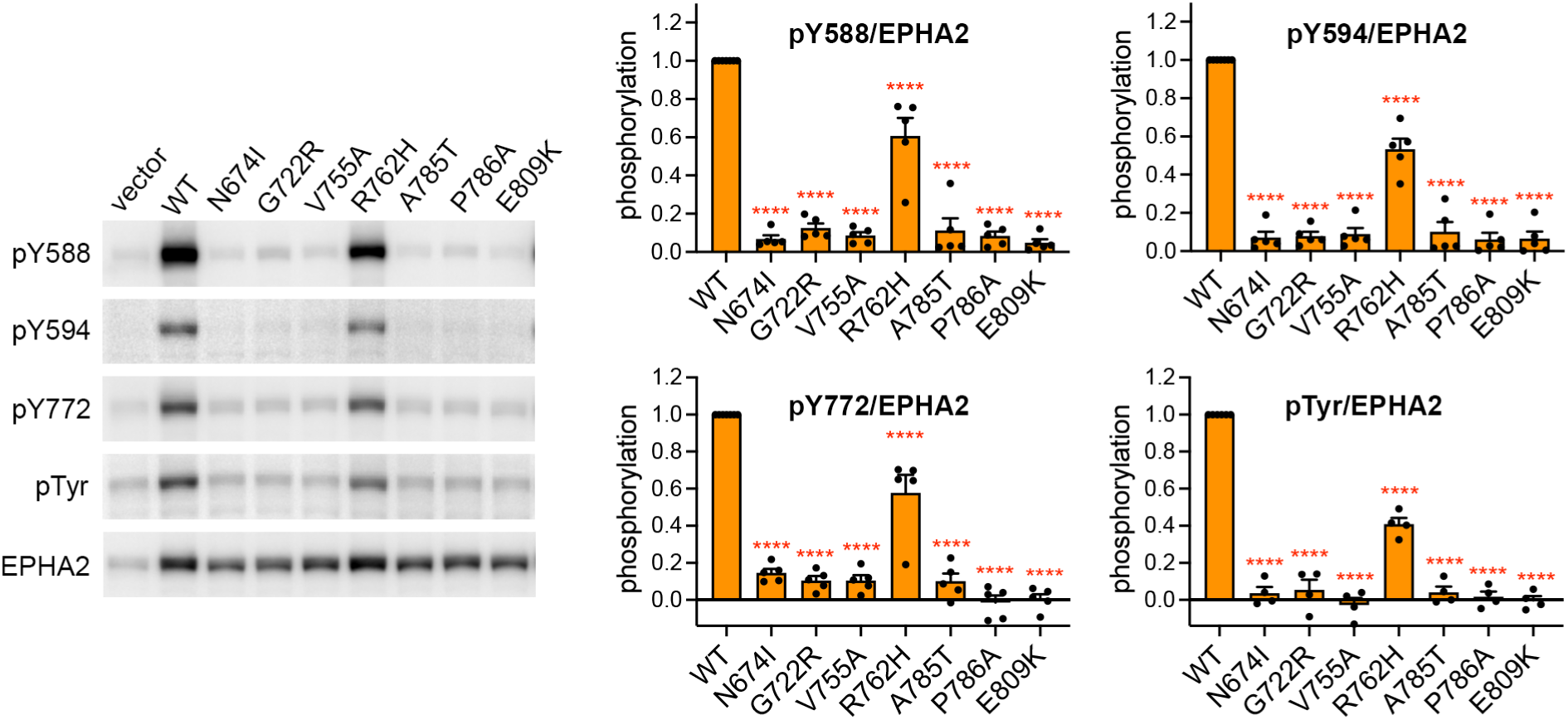
Mutations in the kinase domain abolish or decrease EPHA2 tyrosine phosphorylation Lysates of HEK293 cells transiently transfected with EPHA2 WT and the indicated mutants were probed by immunoblotting for EPHA2 phosphorylation on Y588, Y594, Y772 and overall tyrosine phosphorylation (pTyr) and for EPHA2 levels. The signal in the vector control lane was subtracted from the pY772 and pTyr signals of transfected EPHA2 WT and mutants before normalization to EPHA2 WT in each experiment. The bars in the graphs show averages ± SE from 5-7 experiments. The individual measurements are shown as black dots. The asterisks indicate the significance of the difference from EPHA2 WT, calculated using one-way ANOVA followed by the Dunnett’s multiple comparisons test (****, p < 0.0001).

### Missense mutations in the @ibronectin domains and juxtamembrane segment moderately affect EPHA2 tyrosine phosphorylation

We also assessed the effects on EPHA2 tyrosine phosphorylation of four missense mutations located outside the kinase domain and predicted to affect receptor function (Fig. 4). These mutations include P351S in the first fibronectin domain, Y503C and T526M in the second fibronectin domain, and K586N in the juxtamembrane segment (Fig. 1B and Table S1). Among them, P351, Y503 and T526 are conserved in all Eph receptors (51), suggesting a key role in the activities of the Eph receptor family. We found that the Y503C and T526M mutations decrease overall EPHA2 tyrosine phosphorylation (pTyr) and, in the case of Y503C, also phosphorylation on Y594 and Y772 (Fig. 4).

**Figure 4.**
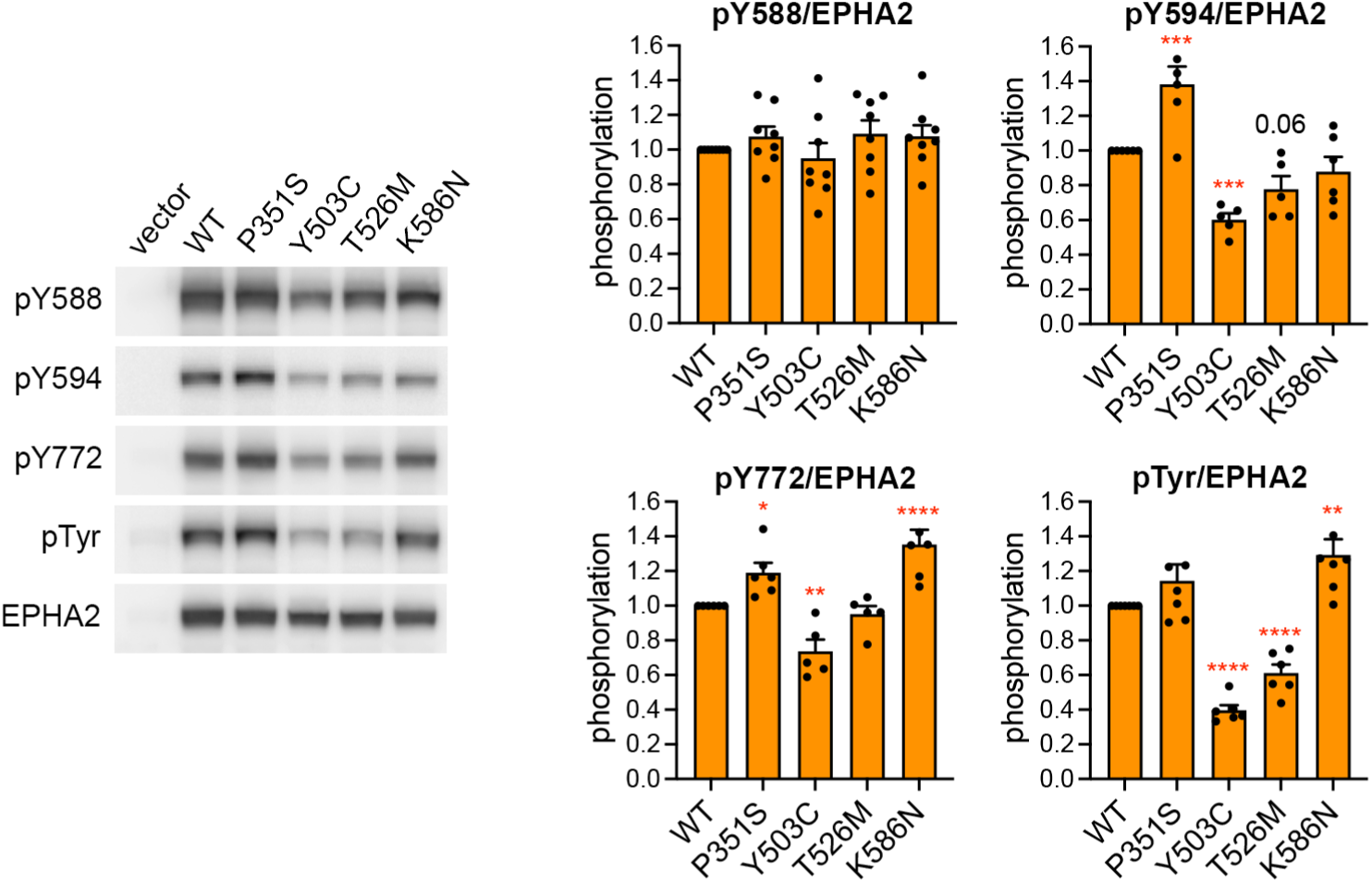
Mutations in the @ibronectin domains and juxtamembrane segment have moderate effects on some EPHA2 tyrosine phosphorylation sites Lysates of HEK293 cells transiently transfected with EPHA2 WT and the indicated mutants were probed by immunoblotting for EPHA2 phosphorylation on Y588, Y594, Y772 and overall tyrosine phosphorylation (pTyr) and for EPHA2 levels. The signal in the vector control lane was subtracted from the phosphorylation signals in transfected EPHA2 WT and mutants before normalization to the signal for EPHA2 WT. The bars in the graphs show averages ± SE from 6-8 experiments. The individual measurements are shown as black dots. The asterisks indicate the significance of the difference from EPHA2 WT, calculated using 2-way ANOVA followed by the Dunnett’s multiple comparisons test (*, p<0.05; **, p < 0.01; ***, p < 0.001; ****, p < 0.0001).

In contrast, we noted a significant increase in EPHA2 P351S phosphorylation on Y594 and Y772. EPHA2 K586N phosphorylation on Y772 and overall tyrosine phosphorylation were also increased. Overall, the four mutations have a less pronounced effect on EPHA2 tyrosine phosphorylation than the mutations in the kinase domain. Furthermore, none of the four mutations seem to significantly change EPHA2 phosphorylation on Y588 (although the K586N mutation may affect the binding affinity of the Y588 phosphospecific antibody). Interestingly, each of the four mutations seems to have distinct consequences on the phosphorylation of different tyrosines.

### Frame-shift and nonsense mutations generate EPHA2 truncated forms lacking all or part of the kinase domain

Most of the EPHA2 mutations in biliary tract tumors are either nonsense or frameshift mutations that introduce a stop codon (Figs. 1A and S2B). Hence, they cause early termination of translation and yield EPHA2 secreted or transmembrane forms that lack the entire kinase domain or its C-terminal portion. We investigated the impact of two EPHA2 frameshift mutations, one representing a highly recurrent mutation that generates a secreted form (P460Rfs*33) and the other representing a mutation that results in a transmembrane form (E623Afs*3). The EPHA2 P460Rfs*33 mutant includes the LBD, Sushi, EGF-like and first fibronectin domain and a few amino acids of the second fibronectin domain, followed by a stretch of 33 amino acids translated in the shifted reading frame (Fig. S3A). The EPHA2 E623Afs*3 mutant includes the entire extracellular region, transmembrane helix, and juxtamembrane segment as well as a few kinase domain amino acids and 3 amino acids translated in the shifted reading frame.

Transient and stable transfections of the EPHA2 P460Rfs*33 construct yielded the expected truncated form secreted in the cell culture medium (Fig. 5A) and also detectable in the cell lysates (Fig. 5B, FLAG blot). The E623Afs*3 construct yielded the expected truncated transmembrane form in the cell lysates (Fig. 5B, FLAG blot). We also observed that the EPHA2 E623Afs*3 mutant, which retains the juxtamembrane tyrosine phosphorylation sites, can be phosphorylated by transiently transfected EPHA2 WT and by endogenous EPHA2 activated by ephrinA1-Fc (Fig. 5B,C). Thus, the EPHA2 E623Afs*3 mutant can serve as a scaffolding protein that binds SH2 domain-containing signaling effectors.

**Figure 5.**
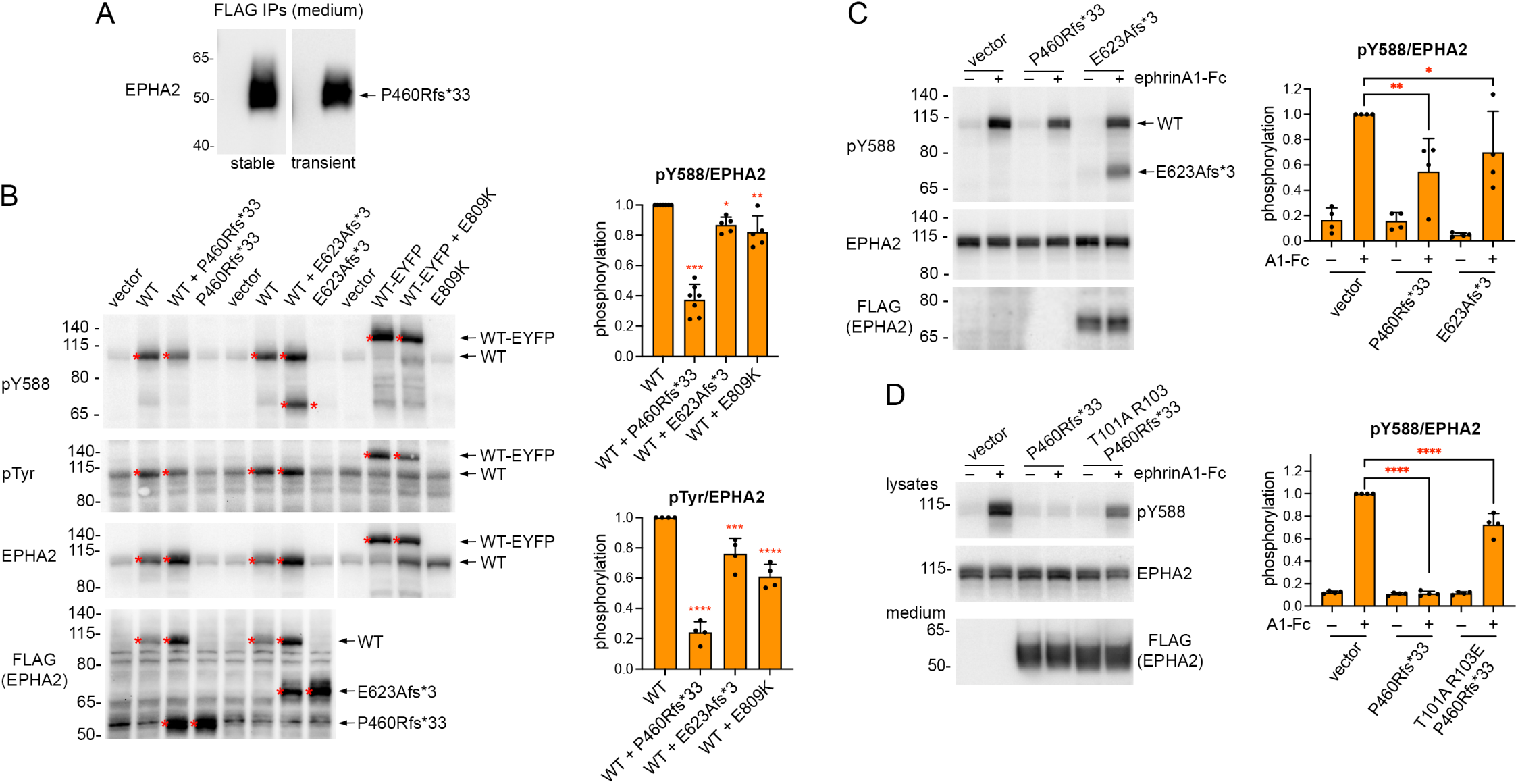
Truncated EPHA2 mutants and a kinase inactive EPHA2 mutant inhibit tyrosine phosphorylation of co-expressed EPHA2 WT. (**A**) Immunoblotting with an antibody to the EPHA2 extracellular region shows the secreted FLAG-EPHA2 P460Rfs*33 fragment in anti-FLAG immunoprecipitates from the conditioned culture medium of stably or transiently transfected HEK293 cells. (**B**) Lysates from HEK293 cells transiently transfected with the indicated constructs were probed by immunoblotting for EphA2 phosphorylation on Y588 (pY588) and overall tyrosine phosphorylation (pTyr). The bars in the graphs show averages ± SE from quantifications of 4-7 experiments, normalized to EPHA2 WT in each experiment. The individual measurements are shown as black dots. The asterisks indicate the significance of the difference from EPHA2 WT, calculated using 1-way ANOVA followed by the Dunnett’s multiple comparisons test for pY588 and 2-way ANOVA followed by the SV ıdák’s multiple comparisons test for pTyr (*, p <0.05; **, p <0.01; ***, p <0.001; ****, p<0.0001). (**C**, **D**) HEK293 cells stably transfected with the indicated mutants were treated for 15 min with 0.5 μg/ml ephrinA1 Fc. Cell lysates were probed by immunoblotting for EPHA2 phosphorylation on Y588 (pY588) and EPHA2. The bars in the graphs show averages ± SE from quantifications of 4 experiments, normalized to endogenous EPHA2 WT in vector control transfected cells stimulated with ephrinA1 Fc. The individual measurements are shown as black dots. The asterisks indicate the significance of the difference from the tyrosine phosphorylation of stimulated endogenous EPHA2 expressed alone, calculated using 2-way ANOVA followed by the SVıdák’s multiple comparisons test (*, p <0.05; **, p<001; ****, p <0.0001). The lower EPHA2 pY588 inhibition by the P460Rfs*33 mutant and the higher variability in C versus D are likely due to the presence of fetal bovine serum in C but not D (see Methods).

### Some EPHA2 mutants have dominant negative effects

Since the EPHA2 P460Rfs*33 mutant contains the LBD and is released into the culture medium, it should inhibit EPHA2 activation by competing with full-length EPHA2 for binding to ephrinA ligands. Indeed, we observed decreased Y588 phosphorylation of endogenous EPHA2 in HEK293 cells stably co-expressing EPHA2 P460Rfs*33 and stimulated with ephrinA1-Fc (Fig. 5C,D). To confirm the mechanism of action responsible for EPHA2 P460Rfs*33-mediated inhibition of endogenous EPHA2 Y588 phosphorylation, we introduced two missense mutations that abrogate ephrin binding in the EPHA2 P460Rfs*33 ephrin-binding pocket (51). We found that the resulting EPHA2 T101A/R103E/P460Rfs*33 triple mutant only partially inhibits ephrinA1-Fc-induced activation of endogenous EPHA2, in contrast to EPHA2 P460Rfs*33 at similar concentration in the medium, which abrogates endogenous EPHA2 Y588 phosphorylation (Fig. 5D). These results show that ephrin ligand sequestration is partially responsible for the dominant negative effects of the EPHA2 P460Rfs*33 mutant. In addition, both P460Rfs*33 and T101A/R103E/P460Rfs*33 mutants may associate with the EPHA2 WT extracellular region (51, 52), decreasing the ability of wild-type receptor molecules to interact with each other and cross-phosphorylate. EPHA2 P460Rfs*33 transiently transfected in HEK293 cells also inhibits Y588 phosphorylation and overall tyrosine phosphorylation of co-transfected EPHA2 WT (Fig. 5B).

The transmembrane E623Afs*3 truncated mutant also contains the LBD but not the kinase domain, and thus would also be expected to exert dominant negative effects. Indeed, when expressed in HEK293 cells (Fig. 5B, FLAG blot), this mutant inhibits Y588 phosphorylation of transiently co-transfected EPHA2 WT (Fig. 5B) as well as ephrinA1-Fc-induced Y588 phosphorylation of endogenous EPHA2 (Fig. 5C).

We also found that a representative kinase-inactive mutant (E809K) exerts dominant negative effects when transiently co-transfected with EPHA2 WT (fused to the EYFP fluorescent protein to increase its size and distinguish it from the E809K mutant in immunoblots; Fig. 5B). As for the other mutants, this is presumably due to competition for ephrin binding and/or oligomerization of the mutant with EPHA2 WT, causing inefficient cross-phosphorylation. These observed dominant negative effects suggest that the EPHA2 truncated and kinase-inactive mutants identified in biliary tract tumors not only are incapable of forward signaling, but can also inhibit signaling by co-expressed EPHA2 WT encoded by the non-mutated allele. Taken together, our findings imply that heterozygous EPHA2 mutations that disrupt kinase activity reduce EPHA2 forward signaling in biliary tract tumors through multiple mechanisms.

### An EPHA2 kinase-inactive mutant induces proliferative masses in a mouse model of cholangiocarcinogenesis

The effects of both truncating and missense mutations strongly suggest a tumor suppressor function for EPHA2 forward signaling (53). Single cell mRNA sequencing data show that cholangiocytes are the cells in the liver with the highest EPHA2 expression and also express high levels of the ephrinA1 ligand (proteinatlas.org). This implies that EPHA2 forward signaling is active in the biliary epithelium and could be disrupted by EPHA2 mutations. To examine the effects of the expression of kinase inactive EPHA2 in the biliary epithelium, we used a validated model of cholangiocarcinogenesis that has been previously used to confirm the *in vivo* role of oncogenes such as YAP1 (17) and tumor suppressors such as FBXW7 (23). We injected transposon constructs encoding EPHA2 WT, the EPHA2 E809K mutant incapable of forward signaling, or the EPHA2 S897A mutant incapable of non-canonical signaling (an oncogenic form of EPHA2 signaling that depends on EPHA2 phosphorylation on S897 (27); see below) into the biliary tree of C57BL/6 mice in combination with a permissive constitutively active myrAKT construct and IL33 administration.

After 12 weeks the mice were euthanized and their livers examined histologically. In the EPHA2 WT and S897A groups, no mice (0/8) developed any tumors. In the E809K group, 25% (2/8) of the mice developed glandular proliferative masses (Fig. 6). These masses are morphologically most consistent with well differentiated CCA, as determined by a dedicated hepatobiliary pathologist, and are positive for the cholangiocyte lineage marker SOX9 (Fig. 6). These results suggest that EPHA2 forward signaling plays a role in suppressing tumor development in the biliary epithelium.

**Figure 6.**
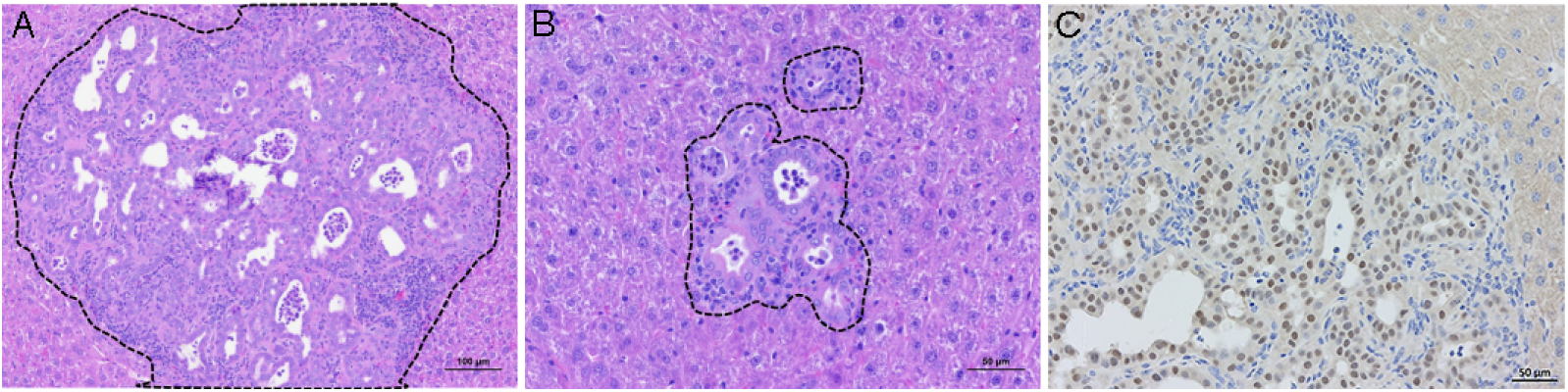
Proliferative lesions induced by biliary instillation of plasmids encoding EPHA2 E809K and myrAKT. (**A,B**) H&E staining of liver sections showing the proliferative lesions observed (black dotted outline) in 2 mice 16 weeks after biliary instillation combined with 3 days of IL33 administration. (**C**) Sox9 staining of the lesion in panel A confirms that the lesion is derived from cells of cholangiocyte lineage. Mouse 1 in A, 10x magnification; mouse 2 in B and mouse 1 in C, 20x magnification.

## DISCUSSION

Biliary tract cancers have an increasing incidence and aggressive biologic behavior and are resistant to systemic, targeted, and immuno therapies (1, 4, 6, 7, 54–56). For most patients, no inciting event or risk factor can be identified, and the mechanisms of tumorigenesis are poorly understood. By analyzing 611 CCA tumor samples with EPHA2 mutation data from 6 studies (including mostly intrahepatic CCA tumors) and 1046 biliary tract tumor samples from the China Pan-cancer study (including intrahepatic and extrahepatic CCA and gallbladder cancer) (48, 57–61) (cbioportal.org), we identified 105 distinct EPHA2 mutations in 122 tumors, corresponding to a 7.4% overall mutation frequency (Tables S1 and S2). This mutation frequency is among the highest for *EPHA2* among all cancer types and is in the same range as other genes whose mutations are under investigation for their role in biliary tract tumor development and progression (cbioportal.org). By comparison, the mutation frequency of well-known mutated genes in the same 7 studies we analyzed is 50% for TP53, 23.5% for KRAS, 17.1% for ARID1A, 12.9% for SMAD4, 8% for BAP1, 7.9% for PBRM1, 7.7% for CDKN2A, 7.1% for PI3KCA, 5.6% for IDH1 and IDH2 combined, and 4.1% for FGFR1-3 combined (cbioportal). Interestingly, the mutation frequency of EPHA2 is somewhat higher in intrahepatic CCA (8.5%) compared to extrahepatic CCA (3.7%) and gallbladder cancer (5.8%) (48). A differential mutation frequency has also been observed for other genes in different subtypes of biliary tract cancer. For example, IDH1/IDH2, PBRM1, FGFR1-3 and BAP1 are preferentially mutated in intrahepatic CCA while KRAS, PRKACA and PRKACB mutations are more prevalent in extrahepatic CCA and TP53, NRAS and BRAF mutations are more prevalent in gallbladder cancer (1, 16, 62–64).

We found that EPHA2 is by far the most highly mutated Eph receptor in biliary tract tumors (Figs. S1 and S2) and that the type and frequency of the mutations are very similar in both sets of studies we analyzed (Tables S1 and S2). Our findings show that most EPHA2 mutations occurring in biliary tract tumors inhibit receptor forward signaling through various mechanisms that depend on the type of mutation. The majority of the EPHA2 mutations are nonsense and frameshift mutations that introduce premature stop codons, leading to the expression of secreted or transmembrane EPHA2 truncated forms lacking the SAM domain and the entire kinase domain or its C-terminal portion. Many of these EPHA2 truncated forms still contain the LBD, which is at the N-terminus, and thus can bind ephrinA ligands but cannot transduce kinase-dependent signals. By analyzing two representative truncated mutants, the secreted P460Rfs*33 and the transmembrane E623Afs*3, as well as the full-length E809K full-length kinase-inactive mutant, we found that these mutations can decrease the tyrosine phosphorylation of co-expressed EPHA2 WT. Through this “dominant negative” effect, EPHA2 heterozygous mutations introducing stop codons or inactivating the kinase domain can not only lead to the loss of kinase activity of EPHA2 encoded by the mutant allele but also decrease the kinase-dependent forward signaling of EPHA2 encoded by the wild-type allele. It will also be interesting to determine whether the unique C-terminal sequences generated by the reading frame shifts could modulate the effects of some EPHA2 truncated mutants. For example, extracellular unpaired cysteines could mediate intermolecular disulfide bonds leading to dimerization of some secreted EPHA2 truncated forms, enabling activation of ephrinA reverse signaling (38). The two cysteines in the unique C-terminal region of the secreted P460Rfs*33 mutant may form an intramolecular disulfide bond, since they do not seem to induce dimerization (as assessed by SDS-PAGE under non-reducing conditions; Fig. S3A,B).

In some instances, the EPHA2 truncated mutants are predicted to contain only a few N-terminal amino acids and thus these mutations essentially cause loss of EPHA2 expression.

Furthermore, some of the premature stop codons may lead to nonsense-mediated mRNA decay (65, 66) and some of the missense mutations may destabilize the EPHA2 protein accelerating its degradation, as has been observed for EPHA2 mutations implicated in cataracts (36). Through these mechanisms, the mutations would decrease EPHA2 expression. In addition, evidence from the TCGA CCA database (cbioportal.org) suggests that loss of EPHA2 heterozygosity may be prevalent in CCA, contributing to decrease EPHA2 expression in a majority of CCA tumors (Fig. S1C) and potentially functioning synergistically with EPHA2 coding mutations.

The EPHA2 missense mutations identified in biliary tract tumors include mutations in the LBD that abrogate ligand binding, and therefore kinase-dependent forward signaling as well as presumably ephrin reverse signaling (28). Two of the mutations involve R103, a residue in the EPHA2 LBD that is critical for ephrin binding (51, 67). On the other hand, the C70W mutation eliminates one of the two disulfide bonds that are important for the structural integrity of the LBD (51, 67). These two different mechanisms likely explain the loss of ligand binding of the LBD mutants. The unpaired cysteine in the EPHA2 LBD of the C70W and R103C mutants mediates dimerization of a portion of the mutant receptor, although the functional consequences of this dimerization remain to be determined (Fig. S3C).

Most mutations in the EPHA2 kinase domain abrogate kinase activity, which is not surprising since all the kinase domain mutations we examined affect residues highly conserved in the Eph receptor family. For example, N674, G722 and V755 are conserved in all Eph receptors except the kinase-inactive EPHA10. R762 in the activation loop is conserved except in the kinase-inactive EPHA10 and EPHB6. A785 is conserved in all Eph receptors except EPHA3 and is part of the “APE” motif in the activation loop, which is highly conserved in kinases and involved in substrate binding, and is thus critical for signaling (53, 68)}. In fact, the APE residues are considered “tumor suppressing hotspot residues” (53). The A785T mutation is also one of the EPHA2 mutations implicated in congenital cataracts, although its effects on EPHA2 kinase activity had not been previously investigated (69, 70). P786 and E809 are conserved in all Eph receptors, and P786 is also part of the APE motif.

The P351S, Y503C, and T526M mutations in the fibronectin domains and the K586N mutation in the juxtamembrane segment show less pronounced effects on EPHA2 phosphorylation than LBD and kinase domain mutations, and some tyrosines in the P351S and K586N mutants show increased rather than decreased phosphorylation. The Y503C mutation in the second fibronectin domain results in an unpaired cysteine that mediates dimerization of a portion of the mutant receptor (Fig. S3C). Interestingly, a number of disease-associated or engineered mutations in other receptor tyrosine kinases, such as the EGF receptor and the fibroblast growth factor receptor, result in unpaired cysteines that induce partial dimerization, with an efficiency that depends on the position of the unpaired cysteine (71–75). The extent of dimer tyrosine phosphorylation/activation in the absence of ligand can be influenced by the spatial arrangement and flexibility of the disulfide linked receptor molecules (72, 73, 76). Consistent with this, we detected tyrosine phosphorylation of the Y503C dimeric mutant but not of the C70W and R103C dimers (Fig. S3C). Interestingly, our data also suggest that the P351S, Y503C, T526M and K586N mutations can differentially affect EPHA2 phosphorylation on different tyrosines. Hence, these mutations may lead to differential activation of EPHA2 downstream signaling pathways, resulting in biased signaling (50, 77).

Fourteen of the EPHA2 mutations were each identified in multiple biliary tract tumors and thus are recurrent (Tables S1 and S2), which is a characteristic of driver mutations. These include the missense mutations involving R103, a residue in the EPHA2 LBD that has been found to be mutated almost exclusively in biliary tract tumors. Here we show that the R103C mutation, which is recurrent in four biliary tract tumors, and the R103P mutation, which is recurrent in two biliary tract tumors, both abolish ephrin binding and ligand-induced EPHA2 phosphorylation. The P460Rfs*33 frameshift mutation is the most frequently detected among all Eph receptor mutations identified in all tumor types (cbioportal) (27). Here we show that the P460Rfs*33 mutation, which was identified in as many as 14 CCA tumors (including one from the UMich PanCancer study in cBioPortal (78), which was not included in our analyses), generates a secreted truncated form of EPHA2 that can inhibit forward signaling by co-expressed EPHA2 WT. The high frequency of the P460Rfs*33 mutation in other digestive system cancers, such as colorectal and gastric cancer (cbioportal.org), is consistent with the notion that disruption of EPHA2 forward signaling may also play a role in driving tumor development outside of the biliary tract. We also found that five of the biliary tract tumors harbor two concurrent EPHA2 mutations, suggesting that both EPHA2 alleles are mutated. Furthermore, in all four EPHA2 mutant TCGA tumors the EPHA2 mutation co-occurs with EPHA2 loss of heterozygosity, suggesting that only mutated EPHA2 is expressed in those tumors. Overall, the pattern of mutations we observed for EPHA2 in biliary tract tumors is characteristic of tumor suppressor genes (53, 79) (intogen.org) and supports a role of EPHA2 as a cancer driver (44, 45).

Our observation that many EPHA2 mutations in biliary tract tumors disrupt forward signaling suggests that ephrinA1-induced EPHA2 signaling is active in the normal biliary epithelium and protects it from tumor initiation. This is in agreement with extensive data in the literature showing that EPHA2 forward signaling can inhibit oncogenic pathways (including AKT/mTORC1, RAS/ERK, and integrin signaling) (24, 27) and can inhibit tumor initiation in pancreatic cancer (80). Consistent with this, in many tumors EPHA2 forward signaling is poorly activated, for example due to low expression of ephrinA ligands in cancer cells with high EPHA2 expression (24, 27, 43, 81) or to the inability of co-expressed EPHA2 and ephrinAs to interact to induce forward signaling (42, 82). Mutations that disrupt ligand binding or kinase activity, such as those found in biliary tract tumors, represent a third mechanism that can limit the tumor suppressing effects of EPHA2 forward signaling on tumorigenesis.

Notably, EPHA2 can also signal through a second form of signaling known as “non-canonical” signaling, which is triggered by phosphorylation of S897 in the kinase-SAM linker region by the AKT, RSK, and PKA kinases (27, 83–85). EPHA2 non-canonical signaling, which is independent of ephrin ligands and kinase activity, can promote cancer cell proliferation, survival, stem-like properties, invasion and metastasis as well as resistance to therapeutic drugs.

Interestingly, EPHA2 forward signaling can inhibit non-canonical signaling. Therefore, EPHA2 missense mutations, and heterozygous truncating mutations occurring in cells that retain expression of the EPHA2 WT allele, may also contribute to tumor progression by enhancing EPHA2 non-canonical signaling.

A previous study indeed reported that an EPHA2 missense mutation identified in lymph node metastases from intrahepatic CCA and known to abolish kinase activity (D739N) (83), promotes tumor growth and metastasis in a mouse model (86). The authors did not investigate the role of kinase activity in the D739N mutant and two other mutants identified in the same study of intrahepatic CCA (I608N and R816W), but focused on the elevated S897 phosphorylation of the mutants compared to EPHA2 WT in CCA cells exposed to ephrinA1-Fc. The pro-oncogenic effects of EPHA2 non-canonical signaling could also explain the reported correlation of CCA tumor growth and metastasis with EPHA2 expression levels in some cases (87). It will be interesting in future studies to more directly evaluate the importance of EPHA2 non-canonical signaling in biliary cancer progression, for example by expressing the EPHA2 S897A mutant deficient in S897 phosphorylation (83–85). It is also conceivable that EPHA1, the only other Eph receptor capable of non-canonical signaling through kinase-SAM linker phosphorylation (88), may compensate for loss of EPHA2 non-canonical signaling in biliary tract tumors expressing EPHA2 truncated forms or lacking EPHA2 expression.

As is typically the case for cancer mutations, EPHA2 mutations likely contribute to the development of biliary tract tumors in concert with other factors. Hence, we examined the *in vivo* effects of the EPHA2 E809K kinase-inactive mutant in a mouse biliary instillation model of CCA development that involves cooperating factors. Our data show that expression of EPHA2 E809K in the biliary tract, but not EPHA2 WT or the EPHA2 S897A mutant, can promote proliferation of normal human cholangiocytes when combined with activated AKT (myrAKT) and administration of the cytokine IL33. This proliferation is associated with development of glandular lesions consistent with well-differentiated CCA.

Activation of AKT, which is not sufficient to cause cholangiocarcinogenesis but functions as a permissive factor in the mouse model we used, is not typically induced by mutations in AKT itself in biliary tract tumors. Rather, mutations in upstream genes can indirectly activate AKT. These include activating mutations in the PI3-kinase subunit PI3KCA and inactivating mutations in the lipid phosphatase PTEN, which have been detected but are rare in biliary tract tumors (1, 16, 59, 63, 64, 91). Increased PI3-kinase/AKT activity in CCA cells can also be induced by downregulation of ARID1A, a subunit of the SWI/SNF chromatin remodeling complex that is considered a tumor suppressor in multiple cancers including CCA (61, 92, 93). Interestingly, ARID1A mutations are frequent in intrahepatic CCA and a high proportion (45%) of the biliary tract tumors with EPHA2 mutations also harbor inactivating ARID1A mutations (adjusted p-value<0.001; cbioportal.org), suggesting a possible interplay between EPHA2 and ARID1A (93). EPHA2 mutations in the China Pan-cancer study also significantly co-occur with mutations in PBRM1, another subunit of the SWI/SNF complex (92), with 20% of the EPHA2 mutant tumors also harboring PBRM1 mutations (adjusted p-value<0.003; cbioportal.org). Furthermore, the EPHA2 gene on chromosome 1p36.13 and the ARID1A gene on chromosome 1p36.11 are a short distance apart, and therefore can be concomitantly affected by chromosomal deletions. In addition, SWI/SNF complexes that include the ARID1A subunit bind to EPHA2 gene regulatory regions to enhance EPHA2 transcription (94).

Because of this, ARID1A loss-of-function can lead to EPHA2 downregulation. Finally, concurrent EPHA2 and ARID1A mutations, but not mutations in EPHA2 or ARID1A individually, significantly correlate with decreased CCA patient survival (93) (Fig. S4), supporting the importance of the EPHA2/ARID1A interplay in CCA aggressiveness and suggesting that concurrent EPHA2 and ARID1A mutations may have diagnostic and prognostic value. Notably, the interplay between EPHA2 and ARID1A/PBRM1 likely has broad significance in cancer, since 33% of the ∼830 tumor samples from all cBioPortal databases that harbor EPHA2 mutations also have ARID1A mutations and 15% also have PBRM1 mutations (adjusted p-value<0.001 for both).

Since inflammation is a known risk factor for the development of biliary tract cancers (1, 3, 6, 17, 18), an interesting speculation is that disruption of epithelial cell junctions in the inflamed epithelium (89) might have unique consequences in the bile tract by facilitating access of the bile acid lithocholic acid to EPHA2. By binding to the ephrin-binding pocket of EPHA2, lithocholic acid can function as an antagonist that inhibits EPHA2/ephrinA1 interaction and downstream forward signaling (90). This could mimic or enhance the effects of EPHA2 mutations.

Although EPHA2 is mutated in ∼2% of all tumors in the cBioPortal databases, the effects of EPHA2 mutations in cancer have not been extensively characterized so far. A study has shown that the T647M and A859D kinase domain mutants, identified in malignant mesothelioma, increased proliferation as compared to EPHA2 WT when expressed in HEK293 cells (95). However, only the kinase-inactive A859D mutant promoted cell migration and reduced responsiveness to the inhibitory effects of the EPHA2 agonist doxazosin, consistent with anti-tumorigenic effects of canonical signaling (95, 96). In contrast, the G391R mutation identified in lung cancer was shown to promote BEAS-2B lung epithelial cell survival and invasion while increasing EPHA2 tyrosine phosphorylation as compared to EPHA2 WT (97). The effects of the G391R mutation may be due at least in part to the loss of an extracellular EPHA2 proteolytic cleavage motif recognized by the MMP14 matrix metalloprotease (81). While EPHA2 mutations in biliary tract tumors – many of which inactivate forward signaling – are increasingly attracting attention (44–46, 86, 93, 98), the information available in the literature on the role of EPHA2 kinase activity in this type of cancer is very limited. A study reported a decrease in EPHA2 tyrosine phosphorylation associated with hypoxia and invasiveness in a CCA cell line (99), whereas another study reported increased EPHA2 tyrosine phosphorylation in a subset of CCA tumors compared to surrounding tissue (100).

Oncogenic effects of EPHA2 forward signaling have been reported in some cases, suggesting an influence of the cellular context and the possibility that cancer cells may in some cases be able to circumvent or hijack EPHA2 kinase activity to promote cancer progression (27). Nevertheless, the potential pro-oncogenic effects due to loss of EPHA2 kinase-dependent forward signaling cautions against the use of EPHA2 kinase inhibitors for the treatment of biliary tract tumors. The information we have obtained on the effects of coding sequence mutations on EPHA2 signaling in biliary tract cancer advance our general understanding of EPHA2 mutations and signaling in cancer (101) and our knowledge of the tumor suppressing kinome (53).

## METHODS

### Databases and analyses of biliary tract tumor samples

In Figs. 1 and S1 and Table S1, we analyzed for Eph receptor mutations 6 of the 10 non-redundant CCA databases available in cBioPortal (cbioportal.org) (102, 103), focusing on the 611 tumor samples profiled for EPHA2 mutations. In Fig. S2 and Table S2, we analyzed EPHA2 mutations in intrahepatic and extrahepatic CCA samples as well as in gallbladder samples from the China Pan-cancer study available in cBioPortal (48). Putative copy number alterations in the CCA TCGA database were determined in cBioPortal according to GISTIC 2.0 (49), with a value of –2 corresponding to homozygous deletion, a value of –1 corresponding to heterozygous loss, and a value of 0 corresponding to no change in gene copy number. Samples with or without mutations were visualized using the “OncoPrint” tool in cBioPortal. The databases were also analyzed for co-occurrence of mutations in EPHA2 and other genes using the “Mutual Exclusivity” analysis in cBioPortal.

### Constructs

N-terminally FLAG-tagged EPHA2 WT in the pLVX-IRES-Neo lentiviral construct and empty pLVX-IRES-Neo lentivirus control were previously described (85). The EPHA2-EYFP constructs in pcDNA3 used for transfection has also been previously described (104). The EPHA2 mutant constructs were generated by overlapping PCR with primers designed to amplify the pLVX-FLAG- EPHA2 WT construct as the template and to introduce the same nucleotide changes identified in EPHA2 mutants from biliary tract tumors. The pLVX-FLAG-EPHA2 T101A/R103E/P460Rfs*33 mutant was generated by overlapping PCR with primers amplifying overlapping portions of the previously generated pLVX-IRES-FLAG-EPHA2 T101A/R103E construct and of the pLVX-FLAG- EPHA2 P460Rfs*33 construct. All amplified portions of the constructs were verified by sequencing.

### Cell culture, transfection and EPHA2 ligand stimulation

The human embryonic kidney cell line HEK293AD was purchased from Cell Biolabs (AD-100). The cells were cultured in Dulbecco’s Modified Eagle Medium (DMEM; Corning, 10–013-CV) supplemented with 10% fetal bovine serum, antibiotics and antimycotics (Corning, 30–004-Cl). The cells were transfected with plasmid constructs using Lipofectamine 2000 (Life Technologies, 11668019). Transiently transfected cells were lysed 24 to 72 hours after transfection. To select stably transfected cells, cells were treated with 1 mg/ml G418 (Life Technologies, 10131035) for ∼9 days after transfection. Cells were periodically tested to verify lack of Mycoplasma contamination and discarded after ∼30 passages.

For ligand stimulations, we used ephrinA1-Fc (R&D Systems, 602-A1-200) and the CGA- WLA-YR custom dimeric peptide purchased from GenScript at >95% purity ((CGAWLAYPDSVPYR)2, dimerized by a disulfide bridge linking Cysteine 1 of the two monomeric precursor peptides) (50). Cells one day after plating (at ∼70% confluency) were treated for 15 min with 0.25μg/ml ephrinA1 Fc (Fig. 2B) or with 4 nM CGA-WLA-YR dimeric peptide (Fig. 2C). Alternatively, cells were plated more sparse and treated two days after platingwith 0.5 μg/ml ephrinA1-Fc for 15 min (Fig. 5C,D). Cells for the experiments in Fig. 5D were also serum starved starting ∼4 hours after plating, which may have increased the dominant negative activity of the secreted EPHA2 P460Rfs*33 mutant.

Lysates were collected in modified RIPA buffer for immunoprecipitation or in LDS-containing sample buffer with 2.5% β-mercaptoethanol for immunoblotting, as described below.

### Immunoprecipitation and pulldown experiments

To obtain lysates for immunoprecipitation, cells were cultured until they reached ∼80% confluence, rinsed once with ice cold PBS containing calcium and magnesium (Lonza, 17–513F), and collected in modified RIPA buffer (1% TX-100, 0.5% sodium deoxycholate, 0.1% SDS, 2 mM EDTA in PBS, pH 7.5) containing Halt Protease and Phosphatase inhibitor cocktail (ThermoFisher Scientific, 78443). Cell lysates were incubated for 5 min on ice with periodic mixing and then centrifuged for 10 min at 16,700 g at 4°C to remove insoluble material. The supernatant was precleared by incubation with Sepharose beads (Sigma-Aldrich, 4B200) for 15 min at 4°C on a rotator followed by centrifugation for 1 min at 4,000 g. Each FLAG immunoprecipitation was performed by incubating 20–25 μl anti-FLAG M2 affinity beads (Sigma-Aldrich, A2220) with precleared cell lysates for 2 hours at 4°C on a rotator. The beads were washed 3 times with 1 ml modified RIPA buffer and once with PBS, and eluted by incubation at 95°C for 2 min in 25 μl LDS-containing sample buffer (Life Technologies, B0007) without β-mercaptoethanol (to avoid dissociating the chains of the FLAG antibody not directly linked to the beads). Following centrifugation for 30 sec at 4,000 g, the supernatant was collected and β-mercaptoethanol was added to a concentration of 2.5%.

For ephrinA1-Fc pulldowns, GammaBind Sepharose beads (GE Healthcare,17-0886-02) were incubated with 2 μg ephrin-A1 Fc (R&D Systems, 602-A1-200) or with 2 μg human Fc (Fisher Scientific, NC9747692) at 4°C for 1 hour. Cells were lysed in 0.5% TX-100 in PBS with Halt Protease and Phosphatase inhibitor cocktail (ThermoFisher Scientific, 78443), and centrifuged and precleared as described above for immunoprecipitations. Each pull down was performed by incubating 20 μl ephrinA1-Fc-coupled GammaBind Sepharose beads with cell lysates for 2 hours at 4°C on a rotator. The beads were then washed 3 times with 1 ml 0.5% TX-100 and once with PBS and eluted by incubation at 95°C for 2 min in 25 μl LDS-containing sample buffer containing 2.5% β- mercaptoethanol.

### Immunoblotting

Cells were rinsed once with ice cold PBS containing calcium and magnesium before being collected in LDS-containing sample buffer (with or without 2.5% β-mercaptoethanol). Conditioned medium was also mixed with LDS sample buffer for immunoblotting. Samples were heated at 95° for 2 min and briefly sonicated. Lysates, conditioned media, immunoprecipitates, and pulldown samples were run on Bolt 4–12% Bis-Tris Plus gels (Invitrogen, NW04125). After semi-dry transfer to PVDF membranes (Immobilon; Sigma-Aldrich, IPVH00010), the membranes were stained with the protein stain amido black (naphthol blue black), incubated for 1 hour at room temperature in blocking buffer (5% bovine serum albumin in TBS containing 0.1% Tween-20), and then incubated in the cold overnight in blocking buffer containing primary antibodies from Cell Signaling Technology to label EPHA2 (6997; 1:3,000 dilution), EPHA2 phosphorylated on Y588 (12677; 1:1,000 dilution), EPHA2 phosphorylated on Y772 (8244, at 1:2,000 dilution), and EPHA2 phosphorylated on Y594 (3970, at 1:2,000 dilution). The monoclonal anti-FLAG M2 antibody (Millipore-Sigma, F1804; 1:1000 dilution) was also used for immunoblotting. After washing, the membranes were incubated with HRP-conjugated secondary antibodies from Thermo Fisher Scientific, including anti-rabbit A16110 at 1:4,000 dilution, anti-mouse A16078 at 1:4000 dilution, and anti-goat A16005 at 1:4000 dilution. For anti-phosphotyrosine blots, PVDF membranes were incubated overnight with a phosphotyrosine antibody conjugated to HRP (Cell Signaling Technology, P-Tyr-100 5465; 1:2,000 dilution). To obtain a chemiluminescence signal, we used the SuperSignal West Dura Extended Duration Substrate kit (Thermo Fisher Scientific, 34076). The signal was captured using the ChemiDoc Touch Imaging System (Bio-Rad), quantified using Image Lab (Bio-Rad), and analyzed using Prism software (GraphPad Software).

### Mouse biliary instillation model of CCA development

To generate a mouse model of cholangiocarcinogenesis mimicking the human disease, we ectopically expressed the EPHA2 E809K mutant, or EPHA2 WT or S897A mutant as controls, in the biliary tract by using the Sleeping Beauty transposon transfection system with concomitant transduction of constitutively active AKT (myrAKT), according to a model used for the YAP1 oncogene and the FBXW7 tumor suppressor (17, 23). Intrabiliary instillation of the transposon/transposase complex was coupled with lobar bile duct ligation in C57BL/6 male mice, followed by intraperitoneal administration of 1 µg IL33 (R&D systems) daily for 3 consecutive days. In this model, typically 95% of the injected mice survive (17).

### Immunohistochemisty

Histological analysis was performed using tissue fixed in 10% formalin and embedded in paraffin. For immunohistochemistry, paraffin-embedded sections were deparaffinized, hydrated and incubated with Sox9 antibody (Cell Signaling Technology, 82630S, 1:600 dilution). The antibody was detected using a Vectastain Elite ABC Universal PLUS kit containing horseradish peroxidase conjugated secondary antibody (Vector Laboratories, PK-8200) and diaminobenzidine (Vector Laboratories, SK-4100). Sections were counterstained with hematoxylin and eosin (H&E) before mounting.

## ACKNOWLEDGMENTS

This work was supported by National Institutes of Health grant R01CA262794 (to EBP) and in part by the Mayo Clinic Hepatobiliary SPORE (NCI/NIH P50 CA210964), Mayo Center for Cell Signaling in Gastroenterology (NIDDK/NIH P30DK084567), and the Mayo Clinic Department of Surgery (to RLS).

## AUTHORS CONTRIBUTIONS

EK performed the experiments shown in Figs. 2-5 and S3, prepared these figures, and co-wrote the manuscript. AN, JWS and DC performed the mouse studies shown in Fig. 6. RG provided expert pathology review of specimens. RLS designed and supervised the mouse studies and reviewed the manuscript. EBP performed the analyses of mutations shown in Figs. 1, S1, S2 and S4, prepared these figures and Tables S1 and S2, supervised the project, and co-wrote the manuscript.

## REFERENCES

1. Brindley, P. J., Bachini, M., Ilyas, S. I., Khan, S. A., Loukas, A., Sirica, A. E., et al. (2021) Cholangiocarcinoma Nat Rev Dis Primers 7, 65 10.1038/s41572-021-00300-2

2. Zhuravleva, E., O’Rourke, C. J., and Andersen, J. B. (2022) Mutational signatures and processes in hepatobiliary cancers Nat Rev Gastroenterol Hepatol 19, 367–382 10.1038/s41575-022-00587-w

3. Ilyas, S. I., Affo, S., Goyal, L., Lamarca, A., Sapisochin, G., Yang, J. D. et al. (2023) Cholangiocarcinoma - novel biological insights and therapeutic strategies Nature reviews Clinical oncology 20, 470–486 10.1038/s41571-023-00770-1

4. Rizvi, S., and Gores, G. J. (2013) Pathogenesis, diagnosis, and management of cholangiocarcinoma Gastroenterology 145, 1215-1229,

5. Baria, K., De Toni, E. N., Yu, B., Jiang, Z., Kabadi, S. M., and Malvezzi, M. (2022) Worldwide Incidence and Mortality of Biliary Tract Cancer Gastro Hep Adv 1, 618–626 10.1016/j.gastha.2022.04.007

6. Banales, J. M., Marin, J. J. G., Lamarca, A., Rodrigues, P. M., Khan, S. A., Roberts, L. R. et al. (2020) Cholangiocarcinoma 2020: the next horizon in mechanisms and management Nat Rev Gastroenterol Hepatol 17, 557–588 10.1038/s41575-020-0310-z

7. Valle, J., Wasan, H., Palmer, D. H., Cunningham, D., Anthoney, A., Maraveyas, A. et al. (2010) Cisplatin plus gemcitabine versus gemcitabine for biliary tract cancer The New England journal of medicine 362, 1273-1281 10.1056/NEJMoa0908721

8. 8. Bang, Y. J., Ueno, M., Malka, D., Chung, H. C., Nagrial, A., Kelley, R. K., et al. (2019) Pembrolizumab (pembro) for advanced biliary adenocarcinoma: Results from the KEYNOTE-028 (KN028) and KEYNOTE-158 (KN158) basket studies J Clin Oncol 37, 4079

9. 9. Oh, D. Y., Ruth He, A., Qin, S., Chen, L. T., Okusaka, T., Vogel, A., et al. (2022) Durvalumab plus Gemcitabine and Cisplatin in Advanced Biliary Tract Cancer NEJM Evid 1, EVIDoa2200015 10.1056/EVIDoa2200015

10. Kelley, R. K., Ueno, M., Yoo, C., Finn, R. S., Furuse, J., Ren, Z. et al. (2023) Pembrolizumab in combination with gemcitabine and cisplatin compared with gemcitabine and cisplatin alone for patients with advanced biliary tract cancer (KEYNOTE-966): a randomised, double-blind, placebo-controlled, phase 3 trial Lancet 401, 1853-1865 10.1016/S0140-6736(23)00727-4

11. Andersen, J. B., Spee, B., Blechacz, B. R., Avital, I., Komuta, M., Barbour, A. et al. (2012) Genomic and genetic characterization of cholangiocarcinoma identifies therapeutic targets for tyrosine kinase inhibitors Gastroenterology 142, 1021-1031 e1015 10.1053/j.gastro.2011.12.005

12. Borad, M. J., Champion, M. D., Egan, J. B., Liang, W. S., Fonseca, R., Bryce, A. H. et al. (2014) Integrated genomic characterization reveals novel, therapeutically relevant drug targets in FGFR and EGFR pathways in sporadic intrahepatic cholangiocarcinoma PLoS Genet 10, e1004135 10.1371/journal.pgen.1004135

13. Graham, R. P., Barr Fritcher, E. G., Pestova, E., Schulz, J., Sitailo, L. A., Vasmatzis, G. et al. (2014) Fibroblast growth factor receptor 2 translocations in intrahepatic cholangiocarcinoma Hum Pathol 45, 1630-1638 10.1016/j.humpath.2014.03.014

14. Farshidfar, F., Zheng, S., Gingras, M. C., Newton, Y., Shih, J., Robertson, A. G. et al. (2017) Integrative Genomic Analysis of Cholangiocarcinoma Identifies Distinct IDH-Mutant Molecular Profiles Cell reports 18, 2780-2794 10.1016/j.celrep.2017.02.033

15. Abou-Alfa, G. K., Sahai, V., Hollebecque, A., Vaccaro, G., Melisi, D., Al-Rajabi, R. et al. (2020) Pemigatinib for previously treated, locally advanced or metastatic cholangiocarcinoma: a multicentre, open-label, phase 2 study The lancet oncology 21, 671–684 10.1016/S1470-2045(20)30109-1

16. Tella, S. H., Kommalapati, A., Borad, M. J., and Mahipal, A. (2020) Second-line therapies in advanced biliary tract cancers The lancet oncology 21, e29–e41 10.1016/S1470-2045(19)30733-8

17. Yamada, D., Rizvi, S., Razumilava, N., Bronk, S. F., Davila, J. I., Champion, M. D. et al. (2015) IL-33 facilitates oncogene-induced cholangiocarcinoma in mice by an interleukin-6-sensitive mechanism Hepatology 61, 1627-1642 10.1002/hep.27687

18. Cadamuro, M., and Strazzabosco, M. (2022) Inflammatory pathways and cholangiocarcinoma risk mechanisms and prevention Advances in cancer research 156, 39–73 10.1016/bs.acr.2022.02.001

19. Kamp, E. J., Dinjens, W. N., Doukas, M., van Marion, R., Verheij, J., Ponsioen, C. Y. et al. (2022) Genetic alterations during the neoplastic cascade towards cholangiocarcinoma in primary sclerosing cholangitis J Pathol 258, 227–235 10.1002/path.5994

20. Miller, L. J., Holmes, I. M., Chen-Yost, H. I., Smola, B., Lew, M., andPang, J. (2024) Detecting Cholangiocarcinoma in the Setting of Primary Sclerosing Cholangitis: Is Biliary Tract Fluorescence In Situ Hybridization Helpful? Cytopathology 10.1111/cyt.13452

21. Miller, L. J., Holmes, I. M., Chen-Yost, H. I., Smola, B., Lew, M., Betz, B. L. et al. (2024) Performance of fluorescence in situ hybridization in biliary brushings with equivocal cytology: an institutional experience J Am Soc Cytopathol 13, 285–290 10.1016/j.jasc.2024.03.002

22. Tomlinson, J. L., Li, B., Yang, J., Loeuillard, E., Stumpf, H. E., Kuipers, H. et al. (2024) Syngeneic murine models with distinct immune microenvironments represent subsets of human intrahepatic cholangiocarcinoma J Hepatol 80, 892–903 10.1016/j.jhep.2024.02.008

23. Wang, J., Wang, H., Peters, M., Ding, N., Ribback, S., Utpatel, K. et al. (2019) Loss of Fbxw7 synergizes with activated Akt signaling to promote c-Myc dependent cholangiocarcinogenesis J Hepatol 71, 742–752 10.1016/j.jhep.2019.05.027

24. Pasquale, E. B. (2010) Eph receptors and ephrins in cancer: bidirectional signalling and beyond Nat Rev Cancer 10, 165–180 10.1038/nrc2806

25. Boyd, A. W., Bartlett, P. F., and Lackmann, M. (2014) Therapeutic targeting of EPH receptors and their ligands Nat Rev Drug Discov 13, 39–62 10.1038/nrd4175

26. Kania, A., and Klein, R. (2016) Mechanisms of ephrin-Eph signalling in development, physiology and disease Nat Rev Mol Cell Biol 17, 240–256 10.1038/nrm.2015.16

27. Pasquale, E. B. (2024) Eph receptors and ephrins in cancer progression Nat Rev Cancer 24, 5–27 10.1038/s41568-023-00634-x

28. Pasquale, E. B. (2005) Eph receptor signalling casts a wide net on cell behaviour Nat Rev Mol Cell Biol 6, 462–475 10.1038/nrm1662

29. 29. Pasquale, E. B. (2024) Eph receptor signaling complexes in the plasma membrane Trends Biochem Sci 10.1016/j.tibs.2024.10.002

30. Walker-Daniels, J., Hess, A. R., Hendrix, M. J., and Kinch, M. S. (2003) Differential regulation of EphA2 in normal and malignant cells Am J Pathol 162, 1037-1042

31. Miao, H., and Wang, B. (2009) Eph/ephrin signaling in epithelial development and homeostasis Int J Biochem Cell Biol 41, 762–770 S1357-2725(08)00278-1 [pii] 10.1016/j.biocel.2008.07.019

32. Wakayama, Y., Miura, K., Sabe, H., and Mochizuki, N. (2011) EphrinA1-EphA2 Signal Induces Compaction and Polarization of Madin-Darby Canine Kidney Cells by Inactivating Ezrin through Negative Regulation of RhoA J Biol Chem 286, 44243–44253 10.1074/jbc.M111.267047

33. Lin, S., Wang, B., and Getsios, S. (2012) Eph/ephrin signaling in epidermal differentiation and disease Semin Cell Dev Biol 23, 92–101 10.1016/j.semcdb.2011.10.017

34. Park, J. E., Son, A. I., and Zhou, R. (2013) Roles of EphA2 in Development and Disease Genes (Basel) 4, 334–357 10.3390/genes4030334

35. Liu, M., Charek, J. G., Vicetti Miguel, R. D., and Cherpes, T. L. (2025) Ephrin-Eph signaling: an important regulator of epithelial integrity and barrier function Tissue Barriers 2462855 10.1080/21688370.2025.2462855

36. 36. Park, J. E., Son, A. I., Hua, R., Wang, L., Zhang, X., andZhou, R. (2012) Human cataract mutations in EPHA2 SAM domain alter receptor stability and function PLoS One 7, e36564 10.1371/journal.pone.0036564

37. Wilson, K., Shiuan, E., and Brantley-Sieders, D. M. (2021) Oncogenic functions and therapeutic targeting of EphA2 in cancer Oncogene 10.1038/s41388-021-01714-8

38. Barquilla, A., and Pasquale, E. B. (2015) Eph receptors and ephrins: therapeutic opportunities Annu Rev Pharmacol Toxicol 55, 465–487 10.1146/annurev-pharmtox-011112-140226

39. Finney, A. C., Funk, S. D., Green, J. M., Yurdagul, A., Jr., Rana, M. A., Pistorius, R. et al. (2017) EphA2 expression regulates inflammation and fibroproliferative remodeling in atherosclerosis Circulation 136, 566–582 10.1161/CIRCULATIONAHA.116.026644

40. Kaushansky, A., Douglass, A. N., Arang, N., Vigdorovich, V., Dambrauskas, N., Kain, H. S. et al. (2015) Malaria parasites target the hepatocyte receptor EphA2 for successful host infection Science 350, 1089-1092 10.1126/science.aad3318

41. Swidergall, M., Solis, N. V., Wang, Z., Phan, Q. T., Marshall, M. E., Lionakis, M. S. et al. (2019) EphA2 Is a Neutrophil Receptor for Candida albicans that Stimulates Antifungal Activity during Oropharyngeal Infection Cell reports 28, 423–433 e425 10.1016/j.celrep.2019.06.020

42. Zelinski, D. P., Zantek, N. D., Stewart, J. C., Irizarry, A. R., and Kinch, M. S. (2001) EphA2 overexpression causes tumorigenesis of mammary epithelial cells Cancer Res 61, 2301-2306,

43. Macrae, M., Neve, R. M., Rodriguez-Viciana, P., Haqq, C., Yeh, J., Chen, C. et al. (2005) A conditional feedback loop regulates Ras activity through EphA2 Cancer Cell 8, 111–118

44. Martinez-Jimenez, F., Muinos, F., Sentis, I., Deu-Pons, J., Reyes-Salazar, I., Arnedo-Pac, C. et al. (2020) A compendium of mutational cancer driver genes Nat Rev Cancer 20, 555–572 10.1038/s41568-020-0290-x

45. Han, Y., Yang, J., Qian, X., Cheng, W. C., Liu, S. H., Hua, X., et al. (2019) DriverML: a machine learning algorithm for identifying driver genes in cancer sequencing studies Nucleic acids research 47, e45 10.1093/nar/gkz096

46. 46. Zhang, Y., Ma, Z., Li, C., Wang, C., Jiang, W., Chang, J., et al. (2022) The genomic landscape of cholangiocarcinoma reveals the disruption of post-transcriptional modifiers Nat Commun 13, 3061 10.1038/s41467-022-30708-7

47. Zhou, Z. J., Ye, Y. H., Hu, Z. Q., Hou, Y. R., Liu, K. X., Sun, R. Q. et al. (2024) Whole-exome sequencing reveals genomic landscape of intrahepatic cholangiocarcinoma and identifies SAV1 as a potential driver Nat Commun 15, 9960 10.1038/s41467-024-54387-8

48. 48. Wu, L., Yao, H., Chen, H., Wang, A., Guo, K., Gou, W., et al. (2022) Landscape of somatic alterations in large-scale solid tumors from an Asian population Nat Commun 13, 4264 10.1038/s41467-022-31780-9

49. Mermel, C. H., Schumacher, S. E., Hill, B., Meyerson, M. L., Beroukhim, R., andGetz, G. (2011) GISTIC2.0 facilitates sensitive and confident localization of the targets of focal somatic copy-number alteration in human cancers Genome Biology 12, R41 10.1186/gb-2011-12-4-r41

50. 50. Gomez-Soler, M., Gehring, M. P., Lechtenberg, B. C., Zapata-Mercado, E., Ruelos, A., Matsumoto, M. W., et al. (2022) Ligands with different dimeric configurations potently activate the EphA2 receptor and reveal its potential for biased signaling iScience 25, 103870 10.1016/j.isci.2022.103870

51. Seiradake, E., Harlos, K., Sutton, G., Aricescu, A. R., andJones, E. Y. (2010) An extracellular steric seeding mechanism for Eph-ephrin signaling platform assembly Nat Struct Mol Biol 17, 398–402 10.1038/nsmb.1782

52. 52. Himanen, J. P., Yermekbayeva, L., Janes, P. W., Walker, J. R., Xu, K., Atapattu, L., et al. (2010) Architecture of Eph receptor clusters Proc Natl Acad Sci U S A 107, 10860-10865 10.1073/pnas.1004148107

53. Hudson, A. M., Stephenson, N. L., Li, C., Trotter, E., Fletcher, A. J., Katona, G. et al. (2018) Truncation- and motif-based pan-cancer analysis reveals tumor-suppressing kinases Sci Signal 11, 10.1126/scisignal.aan6776

54. Razumilava, N., andGores, G. J. (2013) Classification, diagnosis, and management of cholangiocarcinoma Clin Gastroenterol Hepatol 11, 13–21 10.1016/j.cgh.2012.09.009

55. Zhang, X. F., Bagante, F., Chakedis, J., Moris, D., Beal, E. W., Weiss, M. et al. (2017) Perioperative and Long-Term Outcome for Intrahepatic Cholangiocarcinoma: Impact of Major Versus Minor Hepatectomy J Gastrointest Surg 21, 1841-1850 10.1007/s11605-017-3499-6

56. Sasaki, K., Margonis, G. A., Andreatos, N., Bagante, F., Weiss, M., Barbon, C. et al. (2018) Preoperative Risk Score and Prediction of Long-Term Outcomes after Hepatectomy for Intrahepatic Cholangiocarcinoma J Am Coll Surg 226, 393–403 10.1016/j.jamcollsurg.2017.12.011

57. Ong, C. K., Subimerb, C., Pairojkul, C., Wongkham, S., Cutcutache, I., Yu, W. et al. (2012) Exome sequencing of liver fluke-associated cholangiocarcinoma Nat Genet 44, 690–693 10.1038/ng.2273

58. Jiao, Y., Pawlik, T. M., Anders, R. A., Selaru, F. M., Streppel, M. M., Lucas, D. J. et al. (2013) Exome sequencing identifies frequent inactivating mutations in BAP1, ARID1A and PBRM1 in intrahepatic cholangiocarcinomas Nat Genet 45, 1470-1473 10.1038/ng.2813

59. 59. Zou, S., Li, J., Zhou, H., Frech, C., Jiang, X., Chu, J. S., et al. (2014) Mutational landscape of intrahepatic cholangiocarcinoma Nat Commun 5, 5696 10.1038/ncomms6696

60. 60. Sia, D., Losic, B., Moeini, A., Cabellos, L., Hao, K., Revill, K., et al. (2015) Massive parallel sequencing uncovers actionable FGFR2-PPHLN1 fusion and ARAF mutations in intrahepatic cholangiocarcinoma Nat Commun 6, 6087 10.1038/ncomms7087

61. Jusakul, A., Cutcutache, I., Yong, C. H., Lim, J. Q., Huang, M. N., Padmanabhan, N. et al. (2017) Whole-Genome and Epigenomic Landscapes of Etiologically Distinct Subtypes of Cholangiocarcinoma Cancer discovery 7, 1116-1135 10.1158/2159-8290.CD-17-0368

62. Nakamura, H., Arai, Y., Totoki, Y., Shirota, T., Elzawahry, A., Kato, M. et al. (2015) Genomic spectra of biliary tract cancer Nat Genet 47, 1003-1010 10.1038/ng.3375

63. Lamarca, A., Barriuso, J., McNamara, M. G., andValle, J. W. (2020) Molecular targeted therapies: Ready for “prime time” in biliary tract cancer J Hepatol 73, 170–185 10.1016/j.jhep.2020.03.007

64. Tsilimigras, D. I., Stecko, H., Moris, D., and Pawlik, T. M. (2025) Genomic Profiling of Biliary Tract Cancers: Comprehensive Assessment of Anatomic and Geographic Heterogeneity, Co-Alterations and Outcomes J Surg Oncol 10.1002/jso.28081

65. Tan, K., Stupack, D. G., and Wilkinson, M. F. (2022) Nonsense-mediated RNA decay: an emerging modulator of malignancy Nat Rev Cancer 22, 437–451 10.1038/s41568-022-00481-2

66. Lindeboom, R. G., Supek, F., and Lehner, B. (2016) The rules and impact of nonsense-mediated mRNA decay in human cancers Nat Genet 48, 1112-1118 10.1038/ng.3664

67. Himanen, J. P., Goldgur, Y., Miao, H., Myshkin, E., Guo, H., Buck, M. et al. (2009) Ligand recognition by A-class Eph receptors: crystal structures of the EphA2 ligand-binding domain and the EphA2/ephrin-A1 complex EMBO reports 10, 722–728 embor200991 10.1038/embor.2009.91

68. Gogl, G., Kornev, A. P., Remenyi, A., and Taylor, S. S. (2019) Disordered Protein Kinase Regions in Regulation of Kinase Domain Cores Trends Biochem Sci 44, 300–311 10.1016/j.tibs.2018.12.002

69. Kaul, H., Riazuddin, S. A., Shahid, M., Kousar, S., Butt, N. H., Zafar, A. U. et al. (2010) Autosomal recessive congenital cataract linked to EPHA2 in a consanguineous Pakistani family Mol Vis 16, 511–517 58 [pii]

70. Jarwar, P., Sheikh, S. A., Waryah, Y. M., Ujjan, I. U., Riazuddin, S., Waryah, A. M. et al. (2021) Biallelic Variants in EPHA2 Identified in Three Large Inbred Families with Early-Onset Cataract Int J Mol Sci 22, 10.3390/ijms221910655

71. d’Avis, P. Y., Robertson, S. C., Meyer, A. N., Bardwell, W. M., Webster, M. K., and Donoghue, D. J. (1998) Constitutive activation of fibroblast growth factor receptor 3 by mutations responsible for the lethal skeletal dysplasia thanatophoric dysplasia type I Cell Growth Differ 9, 71–78

72. Moriki, T., Maruyama, H., and Maruyama, I. N. (2001) Activation of preformed EGF receptor dimers by ligand-induced rotation of the transmembrane domain Journal of molecular biology 311, 1011-1026 10.1006/jmbi.2001.4923

73. Lu, C., Mi, L. Z., Grey, M. J., Zhu, J., Graef, E., Yokoyama, S. et al. (2010) Structural evidence for loose linkage between ligand binding and kinase activation in the epidermal growth factor receptor Mol Cell Biol 30, 5432-5443 10.1128/MCB.00742-10

74. Greenall, S. A., Donoghue, J. F., Gottardo, N. G., Johns, T. G., and Adams, T. E. (2015) Glioma- specific Domain IV EGFR cysteine mutations promote ligand-induced covalent receptor dimerization and display enhanced sensitivity to dacomitinib in vivo Oncogene 34, 1658-1666 10.1038/onc.2014.106

75. Sarabipour, S., and Hristova, K. (2016) Pathogenic Cysteine Removal Mutations in FGFR Extracellular Domains Stabilize Receptor Dimers and Perturb the TM Dimer Structure Journal of molecular biology 428, 3903-3910 10.1016/j.jmb.2016.08.026

76. Adar, R., Monsonego-Ornan, E., David, P., andYayon, A. (2002) Differential activation of cysteine-substitution mutants of fibroblast growth factor receptor 3 is determined by cysteine localization Journal of Bone & Mineral Research 17, 860–868,

77. Karl, K., Paul, M. D., Pasquale, E. B., andHristova, K. (2020) Ligand bias in receptor tyrosine kinase signaling J Biol Chem 295, 18494–18507 10.1074/jbc.REV120.015190

78. Robinson, D. R., Wu, Y. M., Lonigro, R. J., Vats, P., Cobain, E., Everett, J. et al. (2017) Integrative clinical genomics of metastatic cancer Nature 548, 297–303 10.1038/nature23306

79. Vogelstein, B., Papadopoulos, N., Velculescu, V. E., Zhou, S., Diaz, L. A., Jr., and Kinzler, K. W. (2013) Cancer genome landscapes Science 339, 1546-1558 10.1126/science.1235122

80. Hill, W., Zaragkoulias, A., Salvador-Barbero, B., Parfitt, G. J., Alatsatianos, M., Padilha, A. et al. (2021) EPHA2-dependent outcompetition of KRASG12D mutant cells by wild-type neighbors in the adult pancreas Curr Biol 31, 2550-2560 e2555 10.1016/j.cub.2021.03.094

81. Sugiyama, N., Gucciardo, E., Tatti, O., Varjosalo, M., Hyytiainen, M., Gstaiger, M. et al. (2013) EphA2 cleavage by MT1-MMP triggers single cancer cell invasion via homotypic cell repulsion J Cell Biol 201, 467–484 10.1083/jcb.201205176

82. Zantek, N. D., Azimi, M., Fedor-Chaiken, M., Wang, B. C., Brackenbury, R., and Kinch, M. S. (1999) E-cadherin regulates the function of the EphA2 receptor tyrosine kinase Cell Growth & Differentiation 10, 629–638,

83. Miao, H., Li, D. Q., Mukherjee, A., Guo, H., Petty, A., Cutter, J. et al. (2009) EphA2 mediates ligand-dependent inhibition and ligand-independent promotion of cell migration and invasion via a reciprocal regulatory loop with Akt Cancer Cell 16, 9–20 10.1016/j.ccr.2009.04.009

84. 84. Zhou, Y., Yamada, N., Tanaka, T., Hori, T., Yokoyama, S., Hayakawa, Y., et al. (2015) Crucial roles of RSK in cell motility by catalysing serine phosphorylation of EphA2 Nat Commun 6, 7679 10.1038/ncomms8679

85. Barquilla, A., Lamberto, I., Noberini, R., Heynen-Genel, S., Brill, L. M., and Pasquale, E. B. (2016) Protein kinase A can block EphA2 receptor-mediated cell repulsion by increasing EphA2 S897 phosphorylation Mol Biol Cell 27, 2757-2770 10.1091/mbc.E16-01-0048

86. 86. Sheng, Y., Wei, J., Zhang, Y., Gao, X., Wang, Z., Yang, J., et al. (2019) Mutated EPHA2 is a target for combating lymphatic metastasis in intrahepatic cholangiocarcinoma Int J Cancer 144, 2440- 2452 10.1002/ijc.31979

87. Cui, X. D., Lee, M. J., Kim, J. H., Hao, P. P., Liu, L., Yu, G. R. et al. (2013) Activation of mammalian target of rapamycin complex 1 (mTORC1) and Raf/Pyk2 by growth factor-mediated Eph receptor 2 (EphA2) is required for cholangiocarcinoma growth and metastasis Hepatology 57, 2248-2260 10.1002/hep.26253

88. 88. Matsumoto, M., Gomez-Soler, M., Lombardi, S., Lechtenberg, B. C., andPasquale, E. B. (2024) Missense mutations of the ephrin receptor EPHA1 associated with Alzheimer’s disease disrupt receptor signaling functions J Biol Chem 301, 108099 10.1016/j.jbc.2024.108099

89. Ivanov, A. I., Parkos, C. A., andNusrat, A. (2010) Cytoskeletal regulation of epithelial barrier function during inflammation Am J Pathol 177, 512–524 10.2353/ajpath.2010.100168

90. 90. Giorgio, C., Hassan Mohamed, I., Flammini, L., Barocelli, E., Incerti, M., Lodola, A., et al. (2011) Lithocholic Acid Is an Eph-ephrin Ligand Interfering with Eph-kinase Activation PLoS One 6, e18128 10.1371/journal.pone.0018128

91. Valle, J. W., Lamarca, A., Goyal, L., Barriuso, J., andZhu, A. X. (2017) New Horizons for Precision Medicine in Biliary Tract Cancers Cancer discovery 7, 943–962 10.1158/2159-8290.CD-17-0245

92. Mittal, P., andRoberts, C. W. M. (2020) The SWI/SNF complex in cancer - biology, biomarkers and therapy Nature reviews Clinical oncology 17, 435–448 10.1038/s41571-020-0357-3

93. 93. Tessiri, S., Techasen, A., Kongpetch, S., Namjan, A., Loilome, W., Chan-On, W., et al. (2022) Therapeutic targeting of ARID1A and PI3K/AKT pathway alterations in cholangiocarcinoma PeerJ 10, e12750 10.7717/peerj.12750

94. 94. Raab, J. R., Resnick, S., andMagnuson, T. (2015) Genome-Wide Transcriptional Regulation Mediated by Biochemically Distinct SWI/SNF Complexes PLoS Genet 11, e1005748 10.1371/journal.pgen.1005748

95. 95. Tan, Y. C., Srivastava, S., Won, B. M., Kanteti, R., Arif, Q., Husain, A. N., et al. (2019) EPHA2 mutations with oncogenic characteristics in squamous cell lung cancer and malignant pleural mesothelioma Oncogenesis 8, 49 10.1038/s41389-019-0159-6

96. Petty, A., Idippily, N., Bobba, V., Geldenhuys, W. J., Zhong, B., Su, B. et al. (2018) Design and synthesis of small molecule agonists of EphA2 receptor Eur J Med Chem 143, 1261-1276 10.1016/j.ejmech.2017.10.026

97. Faoro, L., Singleton, P. A., Cervantes, G. M., Lennon, F. E., Choong, N. W., Kanteti, R. et al. (2010) EphA2 mutation in lung squamous cell carcinoma promotes increased cell survival, cell invasion, focal adhesions, and mammalian target of rapamycin activation J Biol Chem 285, 18575–18585 M109.075085 10.1074/jbc.M109.075085

98. Mahipal, A., Tella, S. H., Kommalapati, A., Anaya, D., and Kim, R. (2019) FGFR2 genomic aberrations: Achilles heel in the management of advanced cholangiocarcinoma Cancer Treat Rev 78, 1–7 10.1016/j.ctrv.2019.06.003

99. Vanichapol, T., Leelawat, K., and Hongeng, S. (2015) Hypoxia enhances cholangiocarcinoma invasion through activation of hepatocyte growth factor receptor and the extracellular signal-regulated kinase signaling pathway Mol Med Rep 12, 3265-3272 10.3892/mmr.2015.3865

100. 100. Gu, T. L., Deng, X., Huang, F., Tucker, M., Crosby, K., Rimkunas, V., et al. (2011) Survey of tyrosine kinase signaling reveals ROS kinase fusions in human cholangiocarcinoma PLoS One 6, e15640 10.1371/journal.pone.0015640

101. Wang, G., Xiao, H., Liang, Z., Feng, Y., Wang, L., Feng, Y. et al. (2024) Molecular characteristics and prognostic role of EPHA2 in human tumors via pan-cancer analysis Medicine (Baltimore) 103, e40741 10.1097/MD.0000000000040741

102. Cerami, E., Gao, J., Dogrusoz, U., Gross, B. E., Sumer, S. O., Aksoy, B. A. et al. (2012) The cBio cancer genomics portal: an open platform for exploring multidimensional cancer genomics data Cancer discovery 2, 401–404 10.1158/2159-8290.CD-12-0095

103. 103. Gao, J., Aksoy, B. A., Dogrusoz, U., Dresdner, G., Gross, B., Sumer, S. O., et al. (2013) Integrative analysis of complex cancer genomics and clinical profiles using the cBioPortal Sci Signal 6, pl1 10.1126/scisignal.2004088

104. Singh, D. R., Kanvinde, P., King, C., Pasquale, E. B., andHristova, K. (2018) The EphA2 receptor is activated through induction of distinct, ligand-dependent oligomeric structures Commun Biol 1, 15 10.1038/s42003-018-0017-7

