## Supplementary Information for "Frequent EPHA2 receptor mutations in cholangiocarcinoma disrupt receptor forward signaling supporting a tumor suppressor role"

**Table S1. EPHA2 cholangiocarcinoma mutations**

**Table S2. EPHA2 biliary tract cancer mutations from the China Pan-cancer study**

**Figure S1. EPHA2 is the most mutated Eph receptor in cholangiocarcinoma.** (**A**) Percentage of tumors with mutations in different Eph receptors identified in the 6 cBioPortal cholangiocarcinoma studies in (B). (**B**) "OncoPrints" showing EPHA2 frequencies and types of mutations identified in the 6 combined cholangiocarcinoma studies indicated, including studies by ICGC (61), JHU (58), TCGA, Shangai (59), National University of Singapore (57), and Mount Sinai (60). (**C**) 80% of the 36 TCGA cholangiocarcinoma tumors profiled for mutations and copy number alterations show EPHA2 shallow deletion (predictive of heterozygous loss, HETLOSS) or deep deletion (predictive of homozygous deletion, HOMDEL), with all 4 tumors that harbor an EPHA2 mutation also showing heterozygous loss.

**Figure S2. EPHA2 mutations in patient biliary tract tumors from the China Pan-cancer study.** (**A**) Percentage of tumors with mutations in different Eph receptors identified in biliary tract tumors from the China Pan-cancer study, including intrahepatic and extrahepatic CCA and gallbladder cancer (48). (**B**) "OncoPrints" showing frequencies and types of mutations identified in 555 intrahepatic and 351 extrahepatic CCA tumors and in 240 gallbladder tumors (cbioportal.org). (**C,D**) EPHA2 mutations identified in the 1146 biliary tract tumors (cbioportal.org), which include nonsense mutations (*, red font), frameshift mutations (fs, red font) and splice site mutations (orange font) (B) and missense mutations (C) (see also Table S2). The height of the bars indicates the number of tumors harboring a particular mutation (recurrence). In bold are the mutations also identified in the 6 cBioPortal cholangiocarcinoma studies (Fig. 1; Table S1). Arrows mark the mutations studied (purple, truncating mutations; orange, LBD mutations). The EPHA2 domains are indicated: SP, signal peptide; LBD, ligand-binding domain; EGF, EGF-like domain; Sushi domain; FN, fibronectin type III domain; TM, transmembrane helix; JM, juxtamembrane segment; kinase, kinase domain; SAM, sterile alpha motif domain.

**Figure S3. Effects of mutations that affect the number of cysteine residues on EPHA2 dimerization.** (**A**) The unique C-terminal amino acid sequence of the EPHA2 P460Rfs*33 frameshift mutant contains an arginine replacing proline 460 and 32 additional unique amino acids, including two cysteine residues (shown in red). (**B**) Conditioned culture media from HEK293 cells stably transfected with the EPHA2 P460Rfs*33 mutant were mixed with sample buffer without β-mercaptoethanol or containing β-mercaptoethanol and probed by immunoblotting for EPHA2 using an anti-FLAG antibody. (**C**) Lysates from HEK293 cells transiently transfected with the indicated EPHA2 mutants and containing β-mercaptoethanol or without β-mercaptoethanol were probed by immunoblotting for EPHA2 phosphorylation on Y588, overall tyrosine phosphorylation (pTyr), and EPHA2. The positions of dimers and monomers are indicated.

**Figure S4. Concurrent mutations in EPHA2 and ARID1A correlate with decreased overall cholangiocarcinoma patient survival**. (**A**) Concurrent mutations in EPHA2 and ARID1A correlate with shorter overall patient survival. The analysis includes 15 tumors with both EPHA2 and ARID1A mutations and 174 tumors without EPHA2 or ARID1A mutations. (**B, C**) Mutations in EPHA2 but not ARID1A (B) or ARID1A but not EPHA2 (C) do not significantly correlate with changes in overall patient survival. The analysis in (B) includes 23 tumors with EPHA2 mutations and 212 tumors without EPHA2 mutations. The analysis in (C) includes 53 tumors with ARID1A mutations and 182 tumors without ARID1A mutations Mutation data are from the ICGC study available in cBioPortal (78), which includes 235 samples with both mutation and patient survival data. Median survival times are indicated in the graphs and p values were calculated using the log-rank Mantel–Cox test.

**Table S1. EPHA2 cholangiocarcinoma mutations**

| Study of Origin | Sample ID | Anatomical Subtype | Protein Change | Concurrent Mut [1] or Copy # | # Tumors with Same Mut [2] | Functional Impact [3] | Mutation Type | # Mut in Sample |
|---|---|---|---|---|---|---|---|---|
| Cholangiocarcinoma (ICGC, Cancer Discov 2017) | CCA_JP_208 | Intrahepatic | L13P | | 1 | SIFT: deleterious (0.02); Polyphen-2: possibly_damaging (0.446); AlphaMissense: benign (0.106) | Missense_Mutation | 16 |
| Cholangiocarcinoma (ICGC, Cancer Discov 2017) | CCA_TH_20 | Intrahepatic | A20Rfs*33 | | 1 (2) | | Frame_Shift_Del | 16 |
| Intrahepatic Cholangiocarcinoma (Shanghai, Nat Commun 2014) | ihch_smmu_86 | Intrahepatic | C70W | | 1 | MutationAssessor: high (7.51); SIFT: deleterious (0); Polyphen-2: probably_damaging (0.999); AlphaMissense: pathogenic (0.998) | Missense_Mutation | 43 |
| Cholangiocarcinoma (ICGC, Cancer Discov 2017) | CCA_TH_74 | Intrahepatic | M73Ifs*20 | A785T | 1 | | Frame_Shift_Ins | 8 |
| Cholangiocarcinoma (ICGC, Cancer Discov 2017) | CCA_TH_65 | Intrahepatic | R88* | | 1 | | Nonsense_Mutation | 13 |
| Cholangiocarcinoma (ICGC, Cancer Discov 2017) | CCA_SG_2 | Intrahepatic | G89* | | 1 | | Nonsense_Mutation | 4 |
| Cholangiocarcinoma (ICGC, Cancer Discov 2017) | CCA_TH_22 | Intrahepatic | R93Lfs*71 | | 1 (1) | | Frame_Shift_Ins | 15 |
| Intrahepatic Cholangiocarcinoma (JHU, Nat Genet 2013) | GB19 | Intrahepatic | R103C | | **4** | MutationAssessor: high (7.49); SIFT: deleterious (0); Polyphen-2: probably_damaging (1); AlphaMissense: pathogenic (0.946) | Missense_Mutation | 53 |
| Cholangiocarcinoma (ICGC, Cancer Discov 2017) | CCA_SG_43 | Intrahepatic | R103P | | **2** | MutationAssessor: medium (6.68); SIFT: deleterious (0); Polyphen-2: probably_damaging (1); AlphaMissense: pathogenic (0.997) | Missense_Mutation | 8 |
| Cholangiocarcinoma (ICGC, Cancer Discov 2017) | CCA_JP_186 | Intrahepatic | F139Lfs*23 | | 1 | | Frame_Shift_Del | 5 |
| Cholangiocarcinoma (ICGC, Cancer Discov 2017) | CCA_BR_1 | Intrahepatic | I145Nfs*5 | | **2** | | Frame_Shift_Ins | 3 |
| Cholangiocarcinoma (TCGA, Firehose Legacy) | TCGA-YR-A95A-01 | Perihilar | I145Nfs*5 | ShallowDel | **2** | | Frame_Shift_Ins | 96 |
| Cholangiocarcinoma (ICGC, Cancer Discov 2017) | CCA_SG_23 | Intrahepatic | H210Pfs*183 | | 1 | | Frame_Shift_Del | 8 |
| Cholangiocarcinoma (ICGC, Cancer Discov 2017) | CCA_JP_6 | Intrahepatic | R244Pfs*39 | | 1 | | Frame_Shift_Ins | 4 |
| Cholangiocarcinoma (ICGC, Cancer Discov 2017) | CCA_TH_122 | Intrahepatic | F280Cfs*3 | | 1 | | Frame_Shift_Ins | 19 |
| Cholangiocarcinoma (ICGC, Cancer Discov 2017) | CCA_SG_42 | Perihilar | F314L | S419* | 1 | MutationAssessor: low (3.87); SIFT: deleterious (0); Polyphen-2: benign (0.014); AlphaMissense: pathogenic (0.908) | Missense_Mutation | 74 |
| Cholangiocarcinoma (Nat. Univ. of Singapore, Nat Genet 2012) | R104 | | R327* | | 1 (2) | | Nonsense_Mutation | 33 |
| Cholangiocarcinoma (ICGC, Cancer Discov 2017) | CCA_TH_14 | Intrahepatic | X327_splice | | **2** (5) | | Splice_Site | 14 |
| Intrahepatic Cholangiocarcinoma (JHU, Nat Genet 2013) | CHOL24 | Intrahepatic | S330Pfs*63 | | **2** | | Frame_Shift_Del | 28 |
| Intrahepatic Cholangiocarcinoma (Shanghai, Nat Commun 2014) | ihch_smmu_53 | Intrahepatic | Y334Lfs*47 | | **2** | | Frame_Shift_Ins | 37 |
| Intrahepatic Cholangiocarcinoma (Shanghai, Nat Commun 2014) | ihch_smmu_13 | Intrahepatic | P351S | | 1 (1) | MutationAssessor: medium (5.46); SIFT:deleterious (0); Polyphen-2: probably_damaging (0.999); AlphaMissense: pathogenic (0.960) | Missense_Mutation | 137 |
| Intrahepatic Cholangiocarcinoma (JHU, Nat Genet 2013) | CHOL42 | Intrahepatic | R415Afs*8 | | 2 | | Frame_Shift_Del | 19 |
| Cholangiocarcinoma (ICGC, Cancer Discov 2017) | CCA_JP_89 | Intrahepatic | S419* | | **2** (5) | | Nonsense_Mutation | 4 |
| Cholangiocarcinoma (ICGC, Cancer Discov 2017) | CCA_SG_42 | Perihilar | S419* | F314L | **2** (5) | | Nonsense_Mutation | 74 |
| Cholangiocarcinoma (ICGC, Cancer Discov 2017) | CCA_JP_56 | Perihilar | P460Rfs*33 | T526M | **14** (64) | | Frame_Shift_Del | 54 |
| Cholangiocarcinoma (ICGC, Cancer Discov 2017) | CCA_JP_132 | Intrahepatic | P460Rfs*33 | | **14** (64) | | Frame_Shift_Del | 44 |
| Cholangiocarcinoma (ICGC, Cancer Discov 2017) | CCA_JP_200 | Perihilar | P460Rfs*33 | | **14** (64) | | Frame_Shift_Del | 73 |
| Cholangiocarcinoma (ICGC, Cancer Discov 2017) | CCA_RO_3 | Perihilar | P460Rfs*33 | | **14** (64) | | Frame_Shift_Del | 44 |
| Cholangiocarcinoma (ICGC, Cancer Discov 2017) | CCA_SG_7 | Extrahepatic | P460Rfs*33 | | **14** (64) | | Frame_Shift_Del | 78 |
| Cholangiocarcinoma (ICGC, Cancer Discov 2017) | CCA_TH_13 | Perihilar | P460Rfs*33 | | **14** (64) | | Frame_Shift_Del | 22 |
| Cholangiocarcinoma (TCGA, Firehose Legacy) | TCGA-W5-AA2G-01 | Intrahepatic | K468* | ShallowDel | 1 | | Nonsense_Mutation | 124 |
| Cholangiocarcinoma (ICGC, Cancer Discov 2017) | CCA_JP_121 | Intrahepatic | E470* | | 1 (1) | | Nonsense_Mutation | 45 |
| Cholangiocarcinoma (ICGC, Cancer Discov 2017) | CCA_JP_96 | Intrahepatic | Y473Tfs*20 | | 1 | | Frame_Shift_Del | 2 |
| Cholangiocarcinoma (ICGC, Cancer Discov 2017) | CCA_TH_111 | Perihilar | N480Qfs*116 | | 1 | | Frame_Shift_Ins | 43 |
| Intrahepatic Cholangiocarcinoma (Mount Sinai 2015) | ICC12 | Intrahepatic | Y503C | | 1 | MutationAssessor: high (6.970); SIFT: deleterious (0); Polyphen-2: probably_damaging (1); AlphaMissense: pathogenic (0.996) | Missense_Mutation | 21 |
| Cholangiocarcinoma (ICGC, Cancer Discov 2017) | CCA_JP_61 | Intrahepatic | Q515L | | 1 | MutationAssessor: low (3.30); SIFT: tolerated (0.11); Polyphen-2: benign ( 0.023); AlphaMissense: benign (0.246) | Missense_Mutation | 10 |
| Cholangiocarcinoma (ICGC, Cancer Discov 2017) | CCA_JP_56 | Perihilar | T526M | P460Rfs*33 | 1 (4) | MutationAssessor: medium (6.53); SIFT: deleterious (0); Polyphen-2: probably_damaging (1); AlphaMissense: pathogenic (0.877) | Missense_Mutation | 54 |
| Cholangiocarcinoma (ICGC, Cancer Discov 2017) | CCA_JP_188 | Intrahepatic | R560Efs*34 | | 1 | | Frame_Shift_Del | 1 |
| Cholangiocarcinoma (ICGC, Cancer Discov 2017) | CCA_TH_2 | Perihilar | K586N | | **2** | MutationAssessor: medium (6.40); SIFT: deleterious (0); Polyphen-2: probably_damaging (0.922); AlphaMissense: pathogenic (0.902) | Missense_Mutation | 9 |
| Cholangiocarcinoma (ICGC, Cancer Discov 2017) | CCA_SG_9 | Intrahepatic | E623Afs*3 | | 1 | | Frame_Shift_Del | 10 |
| Cholangiocarcinoma (ICGC, Cancer Discov 2017) | CCA_TH_83 | Perihilar | K638Rfs*11 | | 1 | | Frame_Shift_Del | 8 |
| Cholangiocarcinoma (ICGC, Cancer Discov 2017) | CCA_TH_12 | Perihilar | N674I | | **2** | MutationAssessor: high (7.51); SIFT: deleterious (0); Polyphen-2: probably_damaging (0.999); AlphaMissense:pathogenic (0.987) | Missense_Mutation | 6 |
| Cholangiocarcinoma (TCGA, Firehose Legacy) | TCGA-3X-AAVA-01 | Intrahepatic | N674I | ShallowDel | **2** | MutationAssessor: high (7.51); SIFT: deleterious (0); Polyphen-2: probably_damaging (0.999); AlphaMissense:pathogenic (0.987) | Missense_Mutation | 86 |
| Cholangiocarcinoma (ICGC, Cancer Discov 2017) | CCA_TH_128 | Perihilar | G722R | | 1 | MutationAssessor:high (7.52); SIFT: deleterious (0); Polyphen-2:probably_damaging (1); AlphaMissense: pathogenic (0.986) | Missense_Mutation | 17 |
| Cholangiocarcinoma (ICGC, Cancer Discov 2017) | CCA_JP_117 | Intrahepatic | V755A | | 1 | MutationAssessor: high (7.13); SIFT: deleterious (0); Polyphen-2: probably_damaging (0.997); AlphaMissense: pathogenic (0.989) | Missense_Mutation | 4 |
| Cholangiocarcinoma (ICGC, Cancer Discov 2017) | CCA_JP_124 | Intrahepatic | R762H | | 1 (1) | MutationAssessor:medium (6.19); SIFT: deleterious (0); Polyphen-2: probably_damaging (0.997); AlphaMissense: pathogenic (0.958) | Missense_Mutation | 120 |
| Cholangiocarcinoma (ICGC, Cancer Discov 2017) | CCA_TH_74 | Intrahepatic | A785T | M73Ifs*20 | 1 | MutationAssessor: high (7.49); SIFT: deleterious (0); Polyphen-2: probably_damaging (0.999); AlphaMissense: pathogenic (0.9869) | Missense_Mutation | 8 |
| Cholangiocarcinoma (ICGC, Cancer Discov 2017) | CCA_FR_6 | Intrahepatic | P786A | | 1 | MutationAssessor: high (7.16); SIFT: deleterious (0); Polyphen-2: probably_damaging (0.997); AlphaMissense: pathogenic (0.911) | Missense_Mutation | 6 |
| Cholangiocarcinoma (ICGC, Cancer Discov 2017) | CCA_TH_116 | Intrahepatic | E809K | | 1 | MutationAssessor: high (7.52); SIFT: deleterious (0); Polyphen-2: probably_damaging (1); AlphaMissense: pathogenic (0.999) | Missense_Mutation | 15 |
| Cholangiocarcinoma (ICGC, Cancer Discov 2017) | CCA_JP_177 | Intrahepatic | M926K | | 1 | MutationAssessor: low (3.58); SIFT: tolerated (0.17); Polyphen-2: benign (0.027); AlphaMissense: benign (0.117) | Missense_Mutation | 8 |
| Cholangiocarcinoma (TCGA, Firehose Legacy) | TCGA-W5-AA2I-01 | Intrahepatic | D943V | ShallowDel | 1 (1) | MutationAssessor: medium (5.54); SIFT: deleterious (0); Polyphen-2: probably_damaging (0.999); AlphaMissense: pathogenic (0.910) | Missense_Mutation | 106 |

[1] Mut, mutation.

[2] Number of biliary tract tumors with the indicated mutation from 6 CCA studies (listed in this table) and the China pan-cancer study (listed in Table S2), with recurrent biliary tract tumor mutations in bold and recurrent mutations in other tumor types in parentheses).

[3] The functional effects predicted by 4 different programs are indicated, including the descriptor and, in parentheses, the score. Red indicates highest predicted functional impact, followed by dark orange, light orange and grey (which indicates very low impact).

**Table S2. EPHA2 biliary tract cancer mutations from the China Pan-cancer study**

| Study of Origin | Sample ID | Anatomical Subtype | Protein Change | Concurrent Mut [1] | # Tumors with Same Mut [2] | Functional Impact [3] | Mutation Type | # Mut in Sample |
|---|---|---|---|---|---|---|---|---|
| China Pan-cancer (OrigiMed, Nature 2022) | P-6844 | Extrahepatic Cholangiocarcinoma | *1* | | 1 | | Translation_Start_Site | 10 |
| China Pan-cancer (OrigiMed, Nature 2022) | P-6186 | Intrahepatic Cholangiocarcinoma | E28* | | 1 | | Nonsense_Mutation | 7 |
| China Pan-cancer (OrigiMed, Nature 2022) | P-8467 | Gallbladder Carcinoma | D33V | | 1 | MutationAssessor: high (7.37); SIFT: deleterious (0); Polyphen-2: probably_damaging (0.999); AlphaMissense: pathogenic (0.980) | Missense_Mutation | 4 |
| China Pan-cancer (OrigiMed, Nature 2022) | P-0514 | Extrahepatic Cholangiocarcinoma | A37V | | 1 | MutationAssessor: low (2.68); SIFT:tolerated (0.75); Polyphen-2: benign (0.003); AlphaMissense: benign (0.140) | Missense_Mutation | 32 |
| China Pan-cancer (OrigiMed, Nature 2022) | P-8707 | Intrahepatic Cholangiocarcinoma | C70Y | | 1 (1) | MutationAssessor: high (6.96); SIFT: deleterious (0); Polyphen-2: probably_damaging (0.999); AlphaMissense: pathogenic (0.999) | Missense_Mutation | 5 |
| China Pan-cancer (OrigiMed, Nature 2022) | P-6727 | Intrahepatic Cholangiocarcinoma | E92-F95delinsD | | 1 | | In_Frame_Del | 4 |
| China Pan-cancer (OrigiMed, Nature 2022) | P-0168 | Extrahepatic Cholangiocarcinoma | I94Yfs*4 | | 1 | | Frame_Shift_Ins | 5 |
| China Pan-cancer (OrigiMed, Nature 2022) | P-4725 | Intrahepatic Cholangiocarcinoma | R103C | | **4** | MutationAssessor: high (7.49); SIFT: deleterious (0); Polyphen-2: probably_damaging (1); AlphaMissense: pathogenic(0.946) | Missense_Mutation | 6 |
| China Pan-cancer (OrigiMed, Nature 2022) | P-7629 | Intrahepatic Cholangiocarcinoma | R103C | | **4** | MutationAssessor: high (7.49); SIFT: deleterious (0); Polyphen-2: probably_damaging (1); AlphaMissense: pathogenic (0.946) | Missense_Mutation | 12 |
| China Pan-cancer (OrigiMed, Nature 2022) | P-2503 | Gallbladder Carcinoma | R103C | | **4** | MutationAssessor: high (7.49); SIFT: impact: (0); Polyphen-2: probably_damaging (1); AlphaMissense: pathogenicity: pathogenic (0.946) | Missense_Mutation | 7 |
| China Pan-cancer (OrigiMed, Nature 2022) | P-0136 | Gallbladder Carcinoma | R103G | | 1 (1) | MutationAssessor: high (7.49); SIFT: deleterious (0); Polyphen-2: probably_damaging (0.999); AlphaMissense: pathogenic (0.963) | Missense_Mutation | 4 |
| China Pan-cancer (OrigiMed, Nature 2022) | P-4738 | Gallbladder Carcinoma | R103P | | **2** | MutationAssessor: medium ( 6.68); SIFT: deleterious (0); Polyphen-2: probably_damaging (1); AlphaMissense: pathogenic (0.997) | Missense_Mutation | 3 |
| China Pan-cancer (OrigiMed, Nature 2022) | P-5503 | Extrahepatic Cholangiocarcinoma | G111Wfs*33 | | 1 | | Frame_Shift_Ins | 5 |
| China Pan-cancer (OrigiMed, Nature 2022) | P-0368 | Intrahepatic Cholangiocarcinoma | F119Sfs*41 | | 1 | | Frame_Shift_Del | 2 |
| China Pan-cancer (OrigiMed, Nature 2022) | P-6293 | Gallbladder Carcinoma | A146Cfs*4 | | 1 | | Frame_Shift_Ins | 7 |
| China Pan-cancer (OrigiMed, Nature 2022) | P-7465 | Intrahepatic Cholangiocarcinoma | C188R | | 1 (1) | MutationAssessor: impact: high (7.51); SIFT: deleterious (0); Polyphen-2: probably_damaging (0.999); AlphaMissense: pathogenic (0.9993) | Missense_Mutation | 10 |
| China Pan-cancer (OrigiMed, Nature 2022) | P-6551 | Gallbladder Carcinoma | T225Wfs*170 | | 1 | | Frame_Shift_Ins | 6 |
| China Pan-cancer (OrigiMed, Nature 2022) | P-10020 | Intrahepatic Cholangiocarcinoma | G240Vfs*153 | | 1 (21) | | Frame_Shift_Del | 41 |
| China Pan-cancer (OrigiMed, Nature 2022) | P-6114 | Intrahepatic Cholangiocarcinoma | E241* | | **3** (3) | | Frame_Shift_Ins | 5 |
| China Pan-cancer (OrigiMed, Nature 2022) | P-5104 | Intrahepatic Cholangiocarcinoma | E241* | | **3** (3) | | Frame_Shift_Ins | 4 |
| China Pan-cancer (OrigiMed, Nature 2022) | P-4208 | Gallbladder Carcinoma | E241* | | **3** (3) | | Frame_Shift_Ins | 2 |
| China Pan-cancer (OrigiMed, Nature 2022) | P-5485 | Extrahepatic Cholangiocarcinoma | C262Pfs*20 | | 1 | | Frame_Shift_Del | 53 |
| China Pan-cancer (OrigiMed, Nature 2022) | P-5106 | Intrahepatic Cholangiocarcinoma | F281Lfs*112 | | 1 | | Frame_Shift_Del | 32 |
| China Pan-cancer (OrigiMed, Nature 2022) | P-7427 | Intrahepatic Cholangiocarcinoma | E287* | | 1 | | Frame_Shift_Ins | 1 |
| China Pan-cancer (OrigiMed, Nature 2022) | P-0972 | Intrahepatic Cholangiocarcinoma | C307Wfs*86 | | 1 | | Frame_Shift_Del | 3 |
| China Pan-cancer (OrigiMed, Nature 2022) | P-0801 | Intrahepatic Cholangiocarcinoma | Q318* | | 1 (1) | | Nonsense_Mutation | 7 |
| China Pan-cancer (OrigiMed, Nature 2022) | P-0750 | Intrahepatic Cholangiocarcinoma | C325F | | 1 | MutationAssessor: high (6.82);SIFT: deleterious (0); Polyphen-2: probably_damaging (0.998); AlphaMissense: pathogenic (0.9953) | Missense_Mutation | 5 |
| China Pan-cancer (OrigiMed, Nature 2022) | P-8715 | Extrahepatic Cholangiocarcinoma | X327_splice | | **2** (5) | | Splice_Site | 4 |
| China Pan-cancer (OrigiMed, Nature 2022) | P-4952 | Intrahepatic Cholangiocarcinoma | S330Pfs*63 | | **2** | | Frame_Shift_Del | 5 |
| China Pan-cancer (OrigiMed, Nature 2022) | P-0968 | Gallbladder Carcinoma | Y334Lfs*47 | | **2** | | Frame_Shift_Ins | 8 |
| China Pan-cancer (OrigiMed, Nature 2022) | P-6982 | Extrahepatic Cholangiocarcinoma | V383M | | 1 (1) | MutationAssessor: low (4.54); SIFT: deleterious (0.02); Polyphen-2: benign (0.043); AlphaMissense: benign (0.329) | Missense_Mutation | 243 |
| China Pan-cancer (OrigiMed, Nature 2022) | P-3154 | Extrahepatic Cholangiocarcinoma | R384H | | 1 | MutationAssessor: neutral (1.28); SIFT: tolerated (0.2); Polyphen-2: benign (0.003); AlphaMissense: benign (0.077) | Missense_Mutation | 6 |
| China Pan-cancer (OrigiMed, Nature 2022) | P-6224 | Intrahepatic Cholangiocarcinoma | Y385* | | **2** (1) | | Nonsense_Mutation | 7 |
| China Pan-cancer (OrigiMed, Nature 2022) | P-9646 | Intrahepatic Cholangiocarcinoma | Y385* | | **2** (1) | | Nonsense_Mutation | 8 |
| China Pan-cancer (OrigiMed, Nature 2022) | P-4976 | Intrahepatic Cholangiocarcinoma | Y385Lfs*69 | | 1 | | Frame_Shift_Ins | 9 |
| China Pan-cancer (OrigiMed, Nature 2022) | P-7656 | Extrahepatic Cholangiocarcinoma | Y385Pfs*12 | | 1 | | Frame_Shift_Ins | 6 |
| China Pan-cancer (OrigiMed, Nature 2022) | P-3499 | Gallbladder Carcinoma | L402Afs*7 | | 1 | | Frame_Shift_Ins | 7 |
| China Pan-cancer (OrigiMed, Nature 2022) | P-1387 | Intrahepatic Cholangiocarcinoma | E403* | | 1 | | Nonsense_Mutation | 5 |
| China Pan-cancer (OrigiMed, Nature 2022) | P-4387 | Intrahepatic Cholangiocarcinoma | R415Pfs*39 | | 1 (1) | | Frame_Shift_Ins | 3 |
| China Pan-cancer (OrigiMed, Nature 2022) | P-6069 | Intrahepatic Cholangiocarcinoma | N416K | N416Kfs*36 | 1 | MutationAssessor: high (6.81);SIFT: deleterious (0.01); Polyphen-2:probably_damaging (0.947); AlphaMissense:pathogenic (0.999) | Missense_Mutation | 4 |
| China Pan-cancer (OrigiMed, Nature 2022) | P-6069 | Intrahepatic Cholangiocarcinoma | N416Kfs*36 | N416K | 1 | | Frame_Shift_Del | 4 |
| China Pan-cancer (OrigiMed, Nature 2022) | P-0747 | Intrahepatic Cholangiocarcinoma | K441Qfs*13 | | 1 | | Frame_Shift_Ins | 6 |
| China Pan-cancer (OrigiMed, Nature 2022) | P-8687 | Intrahepatic Cholangiocarcinoma | R447C | X572_splice | 1 (2) | MutationAssessor: medium (6.20); SIFT: deleterious (0.02); Polyphen-2: possibly_damaging (0.813); AlphaMissense: ambiguous (0.351) | Missense_Mutation | 13 |
| China Pan-cancer (OrigiMed, Nature 2022) | P-2178 | Intrahepatic Cholangiocarcinoma | P460Rfs*33 | | **14** (64) | | Frame_Shift_Del | 25 |
| China Pan-cancer (OrigiMed, Nature 2022) | P-3423 | Intrahepatic Cholangiocarcinoma | P460Rfs*33 | | **14** (64) | | Frame_Shift_Del | 25 |
| China Pan-cancer (OrigiMed, Nature 2022) | P-7579 | Intrahepatic Cholangiocarcinoma | P460Rfs*33 | | **14** (64) | | Frame_Shift_Del | 79 |
| China Pan-cancer (OrigiMed, Nature 2022) | P-7792 | Intrahepatic Cholangiocarcinoma | P460Rfs*33 | | **14** (64) | | Frame_Shift_Del | 31 |
| China Pan-cancer (OrigiMed, Nature 2022) | P-8131 | Intrahepatic Cholangiocarcinoma | P460Rfs*33 | | **14** (64) | | Frame_Shift_Del | 38 |
| China Pan-cancer (OrigiMed, Nature 2022) | P-8558 | Intrahepatic Cholangiocarcinoma | P460Rfs*33 | | **14** (64) | | Frame_Shift_Del | 38 |
| China Pan-cancer (OrigiMed, Nature 2022) | P-5722 | Extrahepatic Cholangiocarcinoma | P461Rfs*33 | | **14** (64) | | Frame_Shift_Ins | 13 |
| China Pan-cancer (OrigiMed, Nature 2022) | P-7971 | Intrahepatic Cholangiocarcinoma | R465* | | **2** (2) | | Nonsense_Mutation | 4 |
| China Pan-cancer (OrigiMed, Nature 2022) | P-5811 | Gallbladder Carcinoma | R465* | | **2** (2) | | Nonsense_Mutation | 17 |
| China Pan-cancer (OrigiMed, Nature 2022) | P-3307 | Intrahepatic Cholangiocarcinoma | G477Tfs*15 | | 1 | | Frame_Shift_Del | 8 |
| China Pan-cancer (OrigiMed, Nature 2022) | P-2945 | Intrahepatic Cholangiocarcinoma | S479Cfs*112 | | 1 | | Frame_Shift_Del | 6 |
| China Pan-cancer (OrigiMed, Nature 2022) | P-7794 | Intrahepatic Cholangiocarcinoma | Q508* | | 1 | | Nonsense_Mutation | 10 |
| China Pan-cancer (OrigiMed, Nature 2022) | P-8011 | Intrahepatic Cholangiocarcinoma | E530Dfs*66 | | 1 | | Frame_Shift_Ins | 5 |
| China Pan-cancer (OrigiMed, Nature 2022) | P-7491 | Intrahepatic Cholangiocarcinoma | G539Afs*44 | | 1 | | Frame_Shift_Del | 3 |
| China Pan-cancer (OrigiMed, Nature 2022) | P-6944 | Intrahepatic Cholangiocarcinoma | X561_splice | | 1 (2) | | Splice_Site | 4 |
| China Pan-cancer (OrigiMed, Nature 2022) | P-8687 | Intrahepatic Cholangiocarcinoma | X572_splice | R447C | 1 | | Splice_Site | 13 |
| China Pan-cancer (OrigiMed, Nature 2022) | P-7677 | Intrahepatic Cholangiocarcinoma | Y575Lfs*21 | | 1 | | Frame_Shift_Ins | 3 |
| China Pan-cancer (OrigiMed, Nature 2022) | P-5042 | Extrahepatic Cholangiocarcinoma | K586N | | **2** | MutationAssessor:medium (6.40); SIFT: deleterious (0); Polyphen-2: probably_damaging (0.922); AlphaMissense: (pathogenic (0.902) | Missense_Mutation | 3 |
| China Pan-cancer (OrigiMed, Nature 2022) | P-5380 | Gallbladder Carcinoma | H609Ifs*10 | | 1 | | Frame_Shift_Del | 7 |
| China Pan-cancer (OrigiMed, Nature 2022) | P-7481 | Extrahepatic Cholangiocarcinoma | V613G | | 1 | MutationAssessor: medium ( 6.20); SIFT: deleterious (0); Polyphen-2: possibly_damaging (0.742); AlphaMissense: pathogenic (0.920) | Missense_Mutation | 4 |
| China Pan-cancer (OrigiMed, Nature 2022) | P-4672 | Intrahepatic Cholangiocarcinoma | X622_splice | | **2** (3) | | Splice_Site | 4 |
| China Pan-cancer (OrigiMed, Nature 2022) | P-5020 | Intrahepatic Cholangiocarcinoma | X622_splice | | **2** (3) | | Splice_Site | 6 |
| China Pan-cancer (OrigiMed, Nature 2022) | P-0113 | Intrahepatic Cholangiocarcinoma | V627Gfs*6 | | 1 | | Frame_Shift_Del | 5 |
| China Pan-cancer (OrigiMed, Nature 2022) | P-0289 | Intrahepatic Cholangiocarcinoma | S636L | | 1 | MutationAssessor: low (2.92); SIFT: tolerated (0.27); Polyphen-2: benign (0.131); AlphaMissense: benign (0.072) | Missense_Mutation | 21 |
| China Pan-cancer (OrigiMed, Nature 2022) | P-9250 | Gallbladder Carcinoma | F660Sfs*18 | | 1 | | Frame_Shift_Del | 3 |
| China Pan-cancer (OrigiMed, Nature 2022) | P-7435 | Gallbladder Carcinoma | E693* | | 1 (1) | | Nonsense_Mutation | 29 |
| China Pan-cancer (OrigiMed, Nature 2022) | P-0949 | Intrahepatic Cholangiocarcinoma | X705_splice | | 1 | | Splice_Site | 3 |
| China Pan-cancer (OrigiMed, Nature 2022) | P-5549 | Extrahepatic Cholangiocarcinoma | N732Vfs*73 | | 1 | | Frame_Shift_Del | 25 |
| China Pan-cancer (OrigiMed, Nature 2022) | P-9640 | Intrahepatic Cholangiocarcinoma | V800G | | 1 | MutationAssessor: low (5.00); SIFT:deleterious (0); Polyphen-2: probably_damaging (1); AlphaMissense: pathogenic (0.749) | Missense_Mutation | 5 |
| China Pan-cancer (OrigiMed, Nature 2022) | P-6304 | Intrahepatic Cholangiocarcinoma | S802I | | 1 | MutationAssessor: high (7.52); SIFT: deleterious (0); Polyphen-2: probably_damaging (1); AlphaMissense: pathogenic (0.999) | Missense_Mutation | 4 |
| China Pan-cancer (OrigiMed, Nature 2022) | P-2640 | Intrahepatic Cholangiocarcinoma | E820Gfs*7 | | 1 | | Frame_Shift_Del | 8 |
| China Pan-cancer (OrigiMed, Nature 2022) | P-8487 | Gallbladder Carcinoma | E825K | | 1 (3) | MutationAssessor: low (4.55); SIFT: deleterious (0); Polyphen-2: probably_damaging (0.931); AlphaMissense: pathogenic (0.893) | Missense_Mutation | 15 |
| China Pan-cancer (OrigiMed, Nature 2022) | P-1135 | Intrahepatic Cholangiocarcinoma | K863* | | 1 | | Nonsense_Mutation | 4 |

[1] Mut, mutation.

[2] Number of biliary tract tumors with the indicated mutation from 6 CCA studies (listed in this table) and the China pan-cancer study (listed in Table S2), with recurrent biliary tract tumor mutations in bold and recurrent mutations in other tumor types in parentheses).

[3] The functional effects predicted by 4 different programs are indicated, including the descriptor and, in parentheses, the score. Red indicates highest predicted functional impact, followed by dark orange, light orange and grey (which indicates very low impact).

Figure S1

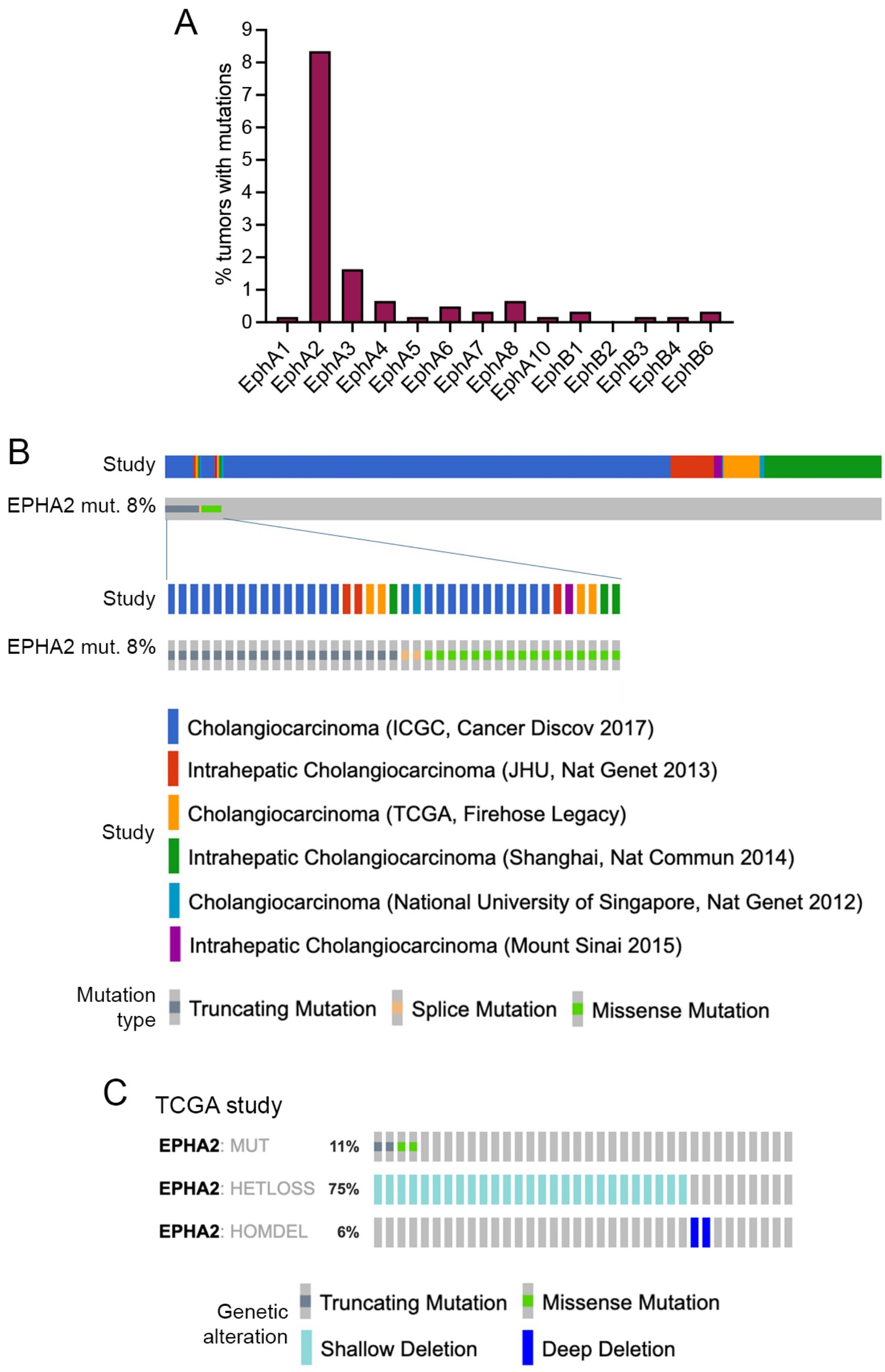
A
% tumors with mutations
0
1
2
3
4
5
6
7
8
9
EphA1
EphA2
EphA3
EphA4
EphA5
EphA6
EphA7
EphA8
EphA10
EphB1
EphB2
EphB3
EphB4
EphB6
B
Study
EPHA2 mut. 8%
Study
EPHA2 mut. 8%
Study
Cholangiocarcinoma (ICGC, Cancer Discov 2017)
Intrahepatic Cholangiocarcinoma (JHU, Nat Genet 2013)
Cholangiocarcinoma (TCGA, Firehose Legacy)
Intrahepatic Cholangiocarcinoma (Shanghai, Nat Commun 2014)
Cholangiocarcinoma (National University of Singapore, Nat Genet 2012)
Intrahepatic Cholangiocarcinoma (Mount Sinai 2015)
Mutation type
Truncating Mutation
Splice Mutation
Missense Mutation
C
TCGA study
EPHA2: MUT 11%
EPHA2: HETLOSS 75%
EPHA2: HOMDEL 6%
Genetic alteration
Truncating Mutation
Missense Mutation
Shallow Deletion
Deep Deletion

Figure S2

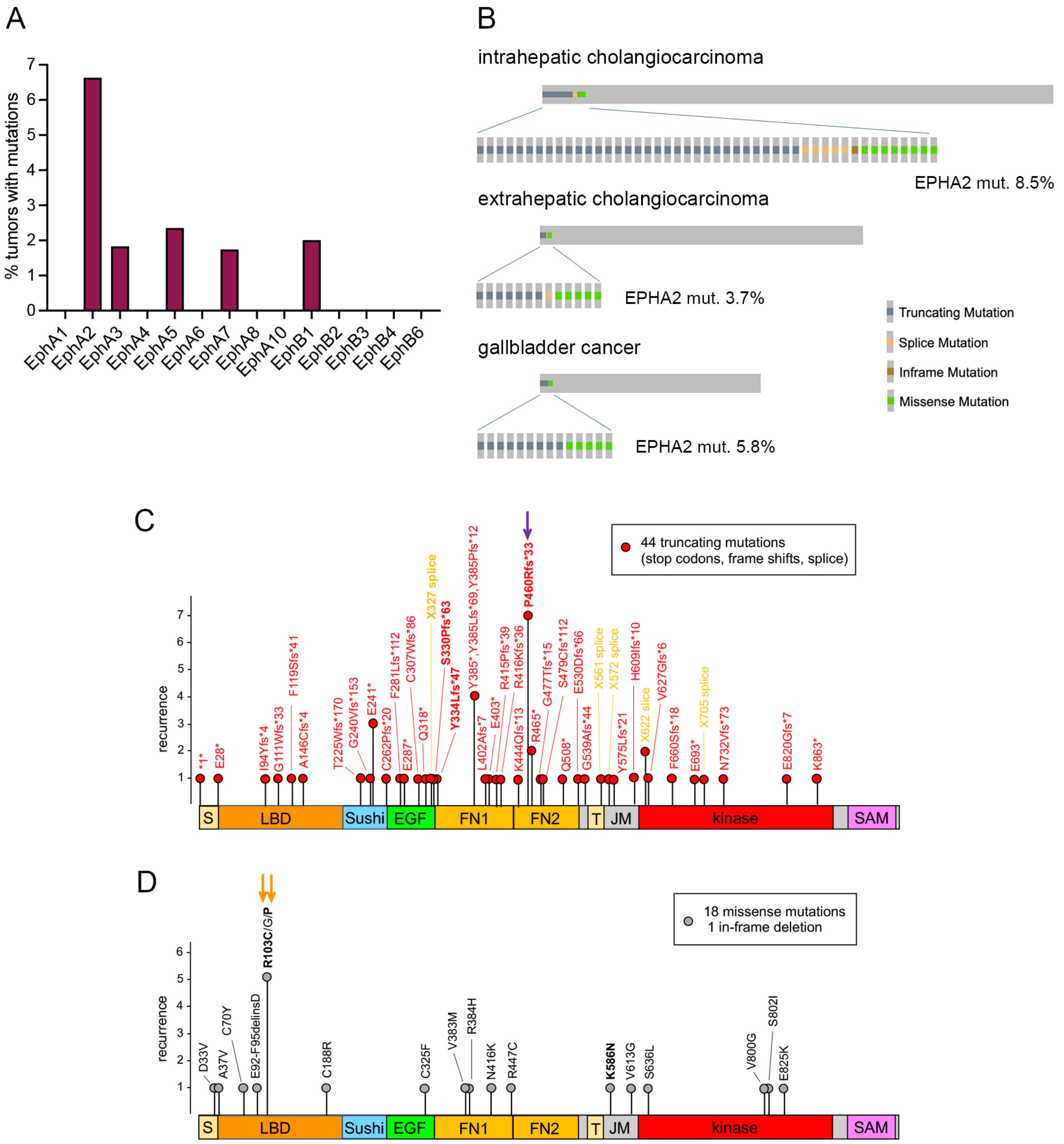
A
% tumors with mutations
EphA1
EphA2
EphA3
EphA4
EphA5
EphA6
EphA7
EphA8
EphA10
EphB1
EphB2
EphB3
EphB4
EphB6
B
intrahepatic cholangiocarcinoma
EPHA2 mut. 8.5%
extrahepatic cholangiocarcinoma
EPHA2 mut. 3.7%
gallbladder cancer
EPHA2 mut. 5.8%
Truncating Mutation
Splice Mutation
Inframe Mutation
Missense Mutation
C
44 truncating mutations
(stop codons, frame shifts, splice)
recurrence
*1*
E28*
I94Yfs*4
G111Wfs*33
F119Sfs*41
A146Cfs*4
T225Wfs*170
G240Vfs*153
E241*
C262Pfs*20
F281Lfs*112
E287*
C307Wfs*86
Q318*
X327 splice
S330Pfs*63
Y334Lfs*47
Y385*,Y385Lfs*69,Y385Pfs*12
L402Afs*7
E403*
R415Pfs*39
R416Kfs*36
K444Qfs*13
P460Rfs*33
R465*
G477Tfs*15
S479Cfs*112
Q508*
E530Dfs*66
G539Afs*44
X561 splice
X572 splice
Y575Lfs*21
H609Ifs*10
X622 splice
V627Gfs*6
F660Sfs*18
E693*
X705 splice
N732Vfs*73
E820Gfs*7
K863*
S
LBD
Sushi
EGF
FN1
FN2
T
JM
kinase
SAM
D
18 missense mutations
1 in-frame deletion
recurrence
D33V
A37V
C70Y
E92-F95delinsD
R103C/G/P
C188R
C325F
V383M
R384H
N416K
R447C
K586N
V613G
S636L
V800G
S802I
E825K

### Figure S3

A

EPHA2 P460Rfs*33 C-terminal unique sequence

RRSRAECGSTRSLTARRETPTATMCAAPRVSP*
460

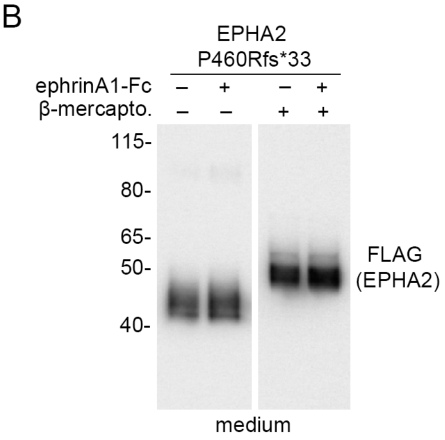


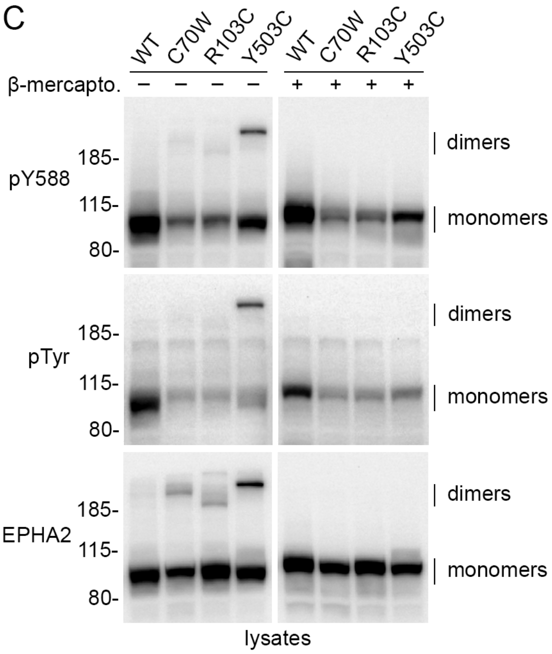

Figure S4

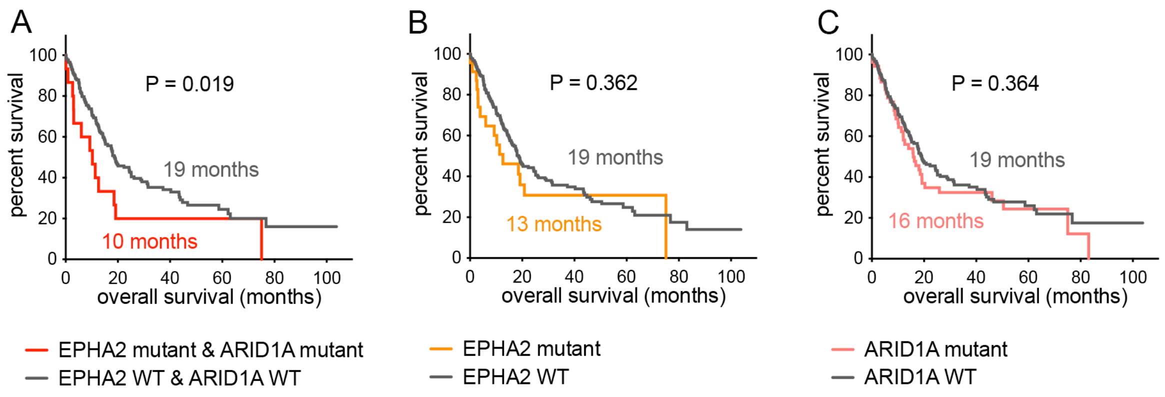

A
P = 0.019
19 months
10 months
percent survival
overall survival (months)
EPHA2 mutant & ARID1A mutant
EPHA2 WT & ARID1A WT
B
P = 0.362
19 months
13 months
percent survival
overall survival (months)
EPHA2 mutant
EPHA2 WT
C
P = 0.364
19 months
16 months
percent survival
overall survival (months)
ARID1A mutant
ARID1A WT
